# Optimised prophylactic vaccination in metapopulations

**DOI:** 10.1101/559732

**Authors:** Mingmei Teo, Nigel Bean, Joshua V. Ross

## Abstract

A highly effective method for controlling the spread of an infectious disease is vaccination. However, there are many situations where vaccines are in limited supply. The ability to determine, under this constraint, a vaccination strategy which minimises the number of people that become infected over the course of a potential epidemic is essential. Two questions naturally arise: when is it best to allocate vaccines, and to whom should they be allocated? We address these questions in the context of metapopulation models of disease spread. We discover that it is optimal to distribute all vaccines prophylactically, rather than withholding until infection is introduced. For small metapopulations, we provide a method for determining the optimal allocation. As the optimal strategy becomes computationally intensive to obtain when the population size increases, we detail an approximation method to determine an approximately optimal vaccination scheme. Through comparisons with other strategies in the literature, we find that our approximate strategy is superior.

## 1 Introduction

Infectious diseases have devastating impacts on society. Examples include the 1918 ‘flu pandemic and the Ebola epidemic in West Africa in 2014. Vaccination is an effective method for controlling the spread of an infectious disease, and in fact eradicated smallpox. However, vaccines are expensive to produce and for some diseases, vaccines may not yet exist and may only be developed during an epidemic. Both of these scenarios result in a limited supply of vaccines. Here we provide new insights to the challenging problem of how best to allocate a limited supply of vaccines, and a new algorithm for use by public health officials.

The optimal allocation of a limited supply of vaccines to a population has been investigated in a range of different contexts, such as metapopulation models [1, 2, 3, 4, 5, 6, 7] and age-structured models [8, 9]. Three main strategies arise from these studies: the *equalising* strategy [1, 2]; the *deterministic* strategy (by Keeling and Shattock [3]); and, a *fair (pro-rata)* strategy [10], used by many as a basis for comparison.

We focus on a stochastic SIR (*susceptible-infectious-recovered*) metapopulation model and prophylactic vaccination, in a susceptible population. We consider a single *attempted* import of infection into the population which may be blocked if infectious contact is with a vaccinated individual. Our objective is to determine a vaccination scheme that minimises the mean final epidemic size (the expected number of people that become infected following an import attempt).

The first question we answer is whether it is better to vaccinate some of the population before the attempted import, or to wait until the location of first import is known. Both of these approaches may have benefits. Vaccinating before infection is present in the population allows for a reduction in the number of susceptible individuals in the population which might reduce the chance of successful import; it also limits the spread of the disease if import is successful. On the other hand, vaccination post-import might allow for vaccine distribution based on the location of the infection(s). We found that for the question of *when* to allocate, it is better to allocate vaccines before the presence of infection in the population.

Next, we explore the problem of *where* to allocate. That is, we consider the optimal pre-allocation of vaccines to the population. As it is computationally intensive to use exact methods to determine the optimal allocation of vaccines for large populations, we develop an approximately optimal allocation. We compare our approximate strategy, as well as the other existing strategies in the literature, with the optimal strategy. We found that our approximate strategy compared well with the optimal strategy and also compared favourably with other strategies from the literature, especially in the case of *medium-sized* populations (for example, overall population of 1800 individuals). Overall, we found that when the optimal strategy cannot be calculated, as is the case when the population size is large, our approximation strategy is preferred and for very large population sizes, a deterministic strategy performs well.

## 2 Model

People live in cities, and animals on farms, for example. This two-level structure can be represented using a metapopulation model where individuals live in *patches* and different rates of transmission can be specified *between* patches and *within* patches. Further, we have also chosen to consider SIR disease dynamics, where each individual in the population is classified as susceptible, infectious or recovered. Under these dynamics, an individual is initially susceptible, and has some chance of becoming infected if contact is made with an infectious individual. After some period of time, the infectious individual recovers and becomes immune to the disease. Note that we have not considered an exposed class, where individuals are infected but cannot transmit the disease, as we are interested in the mean final epidemic size which is independent of the exposed class.

Consider a metapopulation model with *m* patches and SIR disease dynamics. This can be modelled by a continuous-time Markov chain with state space,

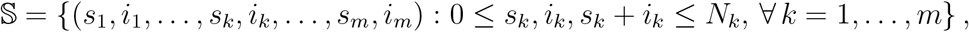

where *N_k_* is the population size of patch *k* and *s_k_* and *i_k_* represent the number of susceptible and infectious individuals in patch *k*, respectively, for *k* = 1, …, *m*. The transition rates for this model, which describe the infection and recovery rates in patch *k*, for *k* = 1,…, *m*, are given by,

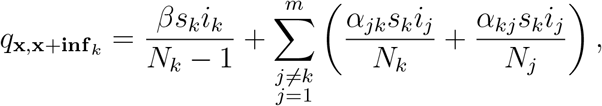

which corresponds to a new infection in patch *k*, and

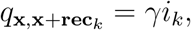

which corresponds to a recovery in patch *k*, where,

- **x** = (*s*_1_,*i*_1_, …, *s_k_*, *i_k_*, …, *s_m_, i_m_*),
- **inf**_*k*_ = **e**_2*k*_ – **e**_2*k*−1_, where **e**_*k*_ is a vector of 0s with a 1 in the *k*^th^ position,
- **rec**_*k*_ = –**e**_2*k*_,
- *γ* is the recovery rate for each infectious individual,
- *β* is the effective transmission rate parameter for transmission within a patch, and
- *α_jk_* is the effective transmission rate parameter for contact between an individual from patch *j* with an individual in patch *k*.

These transition rates form the generator matrix Q for this continuous-time Markov chain. As we are interested in an epidemic that has a positive probability of a major outbreak, we focus on the case *β* > *γ* [11]. Note that we also consider the transmission rates between patches, e.g. *α_jk_*, to be considerably smaller than the transmission rate within a patch, *β*, as transmission within a patch is much more likely than transmission between patches — e.g. transmission into a different city.

Recall that we have chosen to use the mean final epidemic size, which is the expected number of people that become infected over the course of an epidemic, as our measure to minimise. For the SIR model, the mean final epidemic size can also be thought of as the expected number of recovered individuals at the end of the epidemic and can be calculated using path integrals of continuous-time Markov chains [12, 13]. Here, we state the equation used to calculate the mean final epidemic size and refer the reader to [14] for further details. The mean final epidemic size, ζ(**x**), for every state 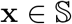, can be obtained by solving,

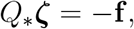

where *Q*_*_ is a modified transition rate matrix of the same size as *Q* and the vector **f** contains the number of recovered individuals for elements corresponding to absorbing states and 0’s elsewhere.

An assumption we make is that the metapopulation is disease-free before attempted import and infection may occur through a single attempted import of infection into the population. To model this, we let *ρ_k_* be the probabiligty that the single attempted import of infection is in patch *k*. This can be represented by 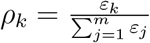, where *ε_k_* is the rate of import into patch k. This form for *ρ_k_* allows for the consideration of different rates of import into different patches, corresponding to different levels of connectivity to the source of infection, say corresponding to air traffic volume with infected countries. If import is attempted at patch *k*, the probability that contact is made with a susceptible individual, and the infection is imported, is 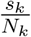. Hence, the probability of an individual in patch *k* becoming infected is 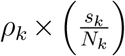, otherwise the infection fails to invade and the final epidemic size is 0. Note that if the entire population is initially susceptible, this probability reduces to *ρ_k_* as *s_k_* = *N_k_*; in this scenario, unsurprisingly, the optimal allocation is identical to the scenario of forced infection, where infection is always assumed to invade (rather than potentially blocked), and this remains true in the case *ε_k_* = *N_k_* because the mean final epidemic size for the latter case is a scaled version of the mean final epidemic size for the former. However, if the rate of import *ε_k_* ≠ *N_k_*, then there is a fundamental difference in the optimal strategy. Figure 1 illustrates this phenomenon.

**Figure 1:**
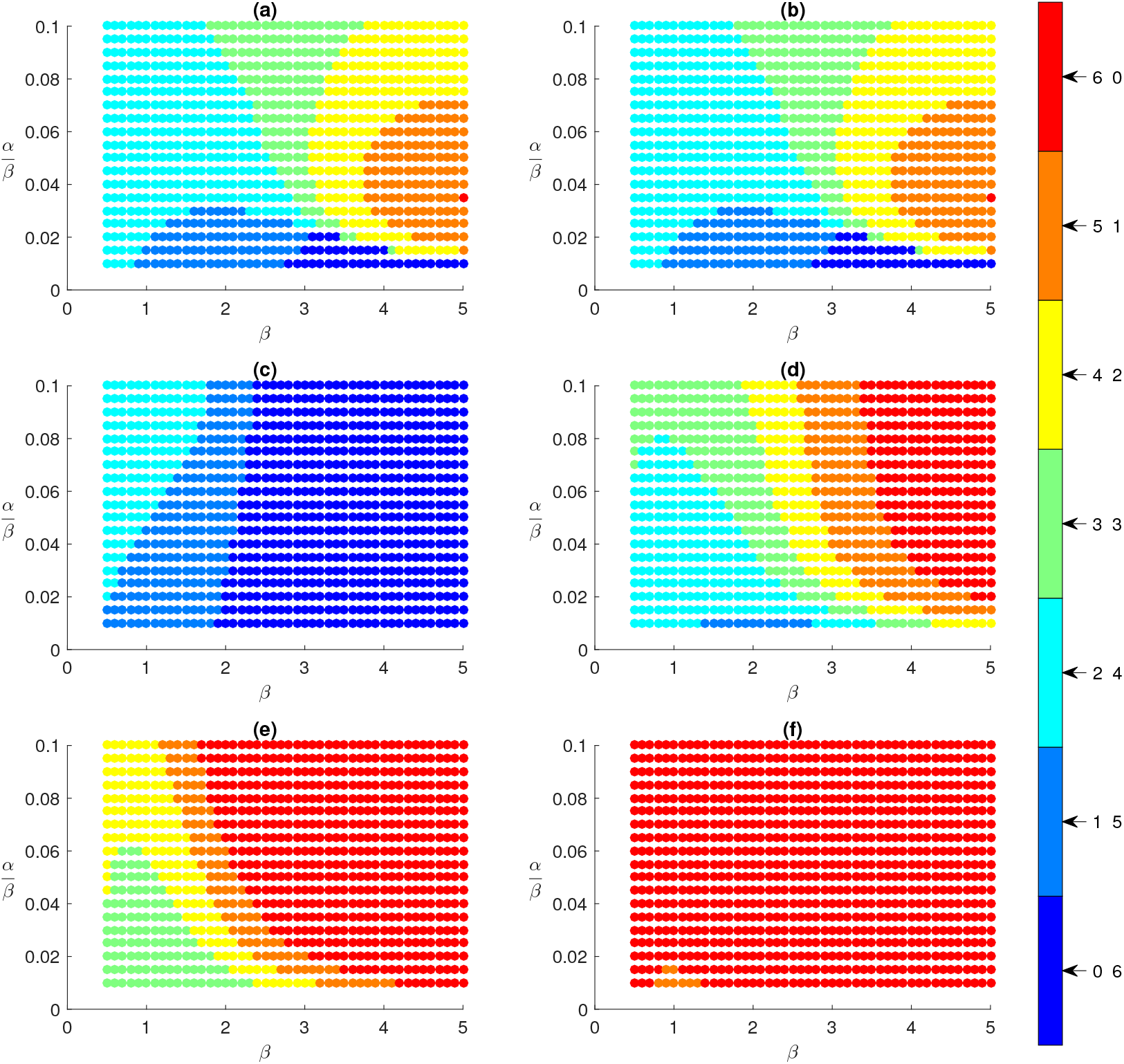
Optimal strategies for a two-patch example with *N*_1_ = 9, *N*_2_ = 15, *V* = 6 vaccines, 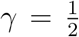, within-patch infection rate *β* ∈ [0.5,5] and cross-patch infection rate *α* ∈ [0.01,0.1] × *β* for different rates of import. (a): Forced infection, (b): Import proportional to population size *ε_k_* = *N_k_* for *k* = 1, 2, (c): *ε*_1_ = 0.35 and *ε*_2_ = 0.65, (d): *ε*_1_ = 0.4 and *ε*_2_ = 0.6, (e): *ε*_1_ = 0.5 and *ε*_2_ = 0.5, (f): *ε*_1_ = 0.75 and *ε*_2_ = 0.25. The colour labels represent the number of vaccines allocated to each patch, (*V*_1_, *V*_2_). This figure shows that the rates of import affect the optimal strategies.

Consider a simple example with two patches of size *N*_1_ = 9 and *N*_2_ = 15, where there are *V* = 6 vaccines. To keep things simple, we have chosen to fix the recovery rate 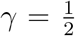 and assume that the cross-patch infection rates are identical, that is, *α*_12_ – *α*_21_ = *α*. We have also chosen to vary the within-patch infection rate *β* ∈ [0.5, 5] and the cross-patch infection rate *a E* [0.01, 0.1] × *β*. Figure 1 consists of the optimal strategies (obtained using the method detailed in Section 4.1) for each (*α, β*) pair and the following rates of import into patch *k, ε_k_*, for *k* = 1, 2,

- *ε_k_* = *N_k_* for *k* =1, 2 (Figure 1b),
- *ε*_1_ = 0.35 and *ε*_2_ = 0.65 (Figure 1c),
- *ε*_1_ = 0.4 and *ε*_2_ = 0.6 (Figure 1d),
- *ε*_1_ = 0.5 and *ε*_2_ = 0.5 (Figure 1e), and
- *ε*_1_ = 0.75 and *ε*_2_ = 0.25 (Figure 1f).

This illustrates the importance of considering attempted import when determining optimal prophylactic vaccination.

Vaccination can be modelled in a variety of ways. For example, vaccinating a susceptible individual can render the individual immune to the disease [1, 2, 3, 4, 5, 6, 7] or vaccinating can reduce an individual’s susceptibility to the disease [15, 16]. Here, we consider *perfect* vaccination, that is, when a susceptible individual is vaccinated, they become immune to the disease. For the SIR model, this can be represented by moving a susceptible individual to the recovered class (and making necessary adjustments to the evaluation of the mean final epidemic size).

## 3 Prophylactic is better than reactive vaccination

First, we explore the question of *when* it is best to allocate vaccines. To do so, we consider three cases:

- allocating all vaccines before infection is present in the population (strict-prophylactic vaccination);
- withholding at least one vaccine until after infection is present in the population (predominantly-prophylactic vaccination); and,
- withholding all vaccines until after infection is present in the population (reactive vaccination).

To compare between these cases and determine the optimal vaccination strategy, we develop a method, the *modified BDP* algorithm (Appendix A.1 Algorithm 1), which utilises Backward Dynamic Programming [17, 18]. The modified BDP algorithm is the standard backward dynamic programming algorithm, with a postprocessing step that accounts for prophylactic vaccination and attempted import of infection. Further, we also incorporate a *delay* between the first onset of infection in the population and vaccination. In a real-world scenario, it is unlikely for public health officials to know instantaneously about the first onset of infection in the population and then be able to apply intervention measures immediately. Hence, there is a delay between these events. To account for this in the modified BDP algorithm, we let *t*_1_ be the time until vaccination can occur after infection is present in the population, measured in days. Then, the transition probabilities of moving from one state to another in time *t*_1_ can be calculated by *P*_1_: = *P*(*t*_1_) = *e*^*Qt*_1_^. In the example considered in Section 3.1, we vary the delay between first infection and vaccination, *t*_1_, to explore how prophylactic vaccination compares with vaccinating after infection is present in the population. For a more detailed explanation of the standard backward dynamic programming algorithm and how it relates to our problem, we refer the reader to Appendix A.1.

Another important aspect of the modified BDP algorithm is how vaccines are allocated during an epidemic. As backward dynamic programming determines the best action to take at each time step, we simplify the modelling, without loss of generality, and limit the number of possible actions by only allowing a single vaccine to be allocated at each time step. Then, given the current state of the process, **S**_*t*_, and the chosen action, *a_t_*, the process moves to the next state **s**′ according to the probability *P*(s′|**S**_*t*_, *a_t_*), which can be obtained from the transition probability matrix, *P*(*t*′) = *e^Qt^*′.

A key point to note here is that a time step, *t*′, does not necessarily correspond to a single day. Hence, the way we have chosen to model the allocation of vaccines during an epidemic does not limit us to allocate at most one vaccine on each day. If we assume that a day corresponds to a time unit of 1, then by choosing values of *t*′ smaller than 1, the process evolves over smaller time steps (than a day) and so vaccines are allocated at a faster rate. For our example, we have chosen to let 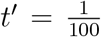 — this essentially means that vaccines are able to be allocated almost simultaneously.

Next, we consider a simple example and apply the modified BDP algorithm to compare between prophylactic vaccination and reactive vaccination.

### 3.1 Results

Consider an example with 40 individuals split between two patches, and there are *V* = 10 vaccines available to be allocated to the population over a fixed time-horizon of *T* = 12 days. For this problem, we have chosen to fix 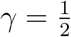 and *β* = 1. Further, we also assume that the cross-patch infection rates, *α_kj_* for *j* ≠ *k* and *k,j* = 1, 2, are identical and set to *α*_12_ = *α*_21_ = *α* = 0.01. Under this set up, we consider four ways to split the population of 40 individuals into two patches of size *N*_1_ and *N*_2_:

1. [*N*_1_, *N*_2_] = [2, 38],
2. [*N*_1_, *N*_2_] = [10, 30],
3. [*N*_1_, *N*_2_] = [15, 25], and
4. [*N*_1_, *N*_2_] = [20, 20].

Then, for each [*N*_1_, *N*_2_], we vary the time until vaccination can occur after infection is present in the population, 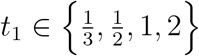, which corresponds to a delay of eight hours, twelve hours, one and two days, respectively.

We apply the modified BDP algorithm (Appendix A.1 Algorithm 1) to each problem to obtain the minimum mean final epidemic size for the vaccination schemes of interest: strict-prophylactic vaccination; predominantly-prophylactic vaccination; and, reactive vaccination. Figure 2 displays the results obtained from the modified BDP algorithm for each problem and each delay *t*_1_.

**Figure 2:**
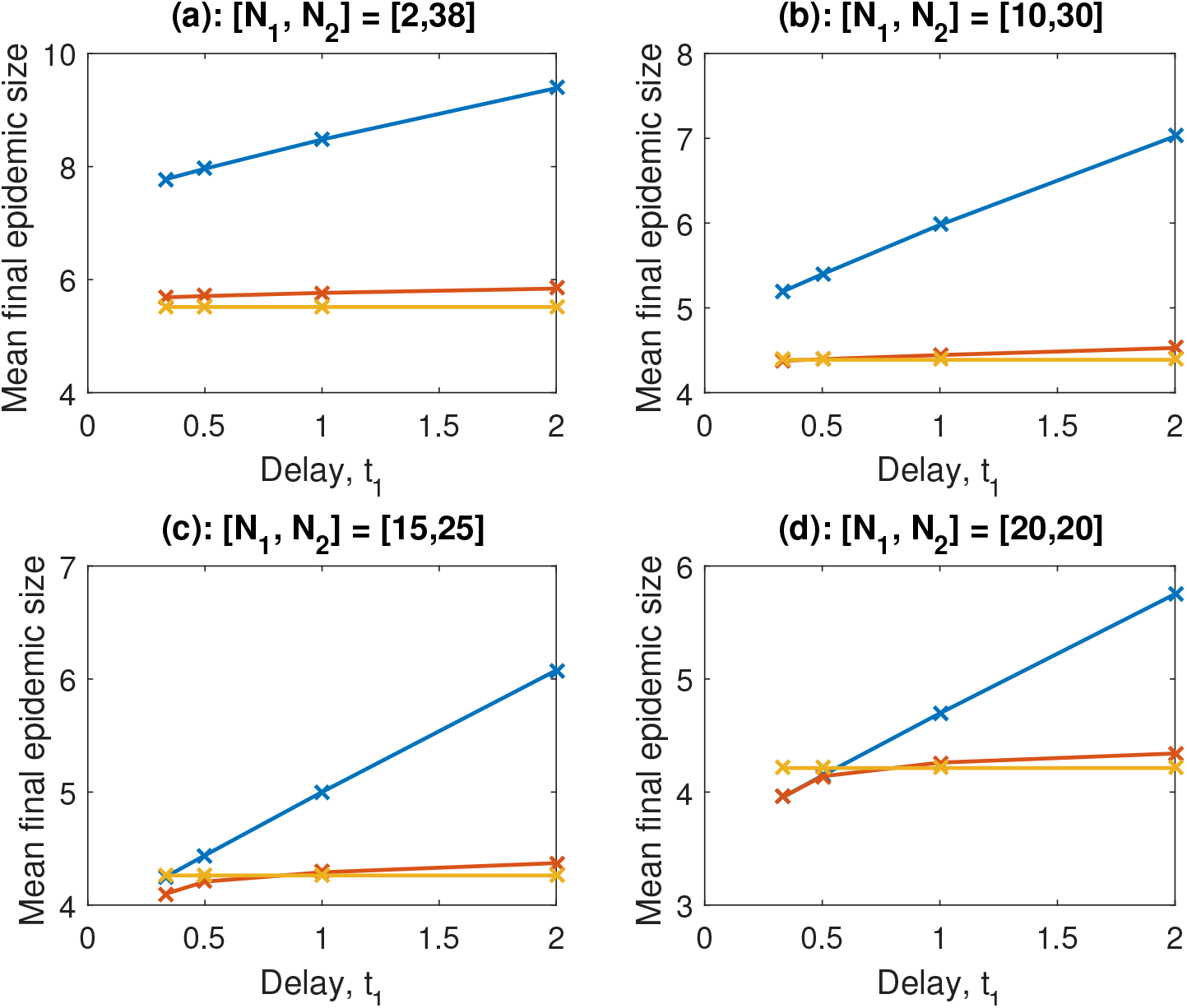
Mean final epidemic size comparison of the three types of vaccination schemes, reactive vaccination (blue), predominantly-prophylactic vaccination (red) and strict-prophylactic vaccination (yellow), for a total population size of 40 individuals and *V* = 10 vaccines by varying *t*_1_, the delay between the first onset of infection in the population and vaccination. The following parameters are assumed fixed: the recovery rate 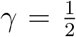, the within-patch infection rate *β* = 1, the cross-patch infection rate *α* = 0.01 and the time-horizon *T* = 12 days. The subplots correspond to different patch sizes, (a): [*N*_1_,*N*_2_] = [2, 38], (b): [*N*_1_,*N*_2_] = [10, 30], (c): [*N*_1_,*N*_2_] = [15, 25], (d): [*N*_1_,*N*_2_] = [20, 20]. Note that the y-axis does not start at zero, but instead has been chosen to make the signal more visible. This shows that, in general, strict-prophylactic vaccination is preferred as it results in smaller mean final epidemic sizes than withholding vaccines until after infection is present in the population, especially with realistic delays.

From Figure 2, we see that, in general, withholding vaccines until after infection is present in the population results in a larger mean final epidemic size than strict-prophylactic vaccination. Hence, in general, strict-prophylactic vaccination is optimal. The only instances where there might be some benefit in withholding vaccines (some or all) until after infection is present in the population occurs when the delay between infection and vaccination is less than 1 day and when the patch sizes are relatively even. The same analysis was performed for different values of within-patch, *β*, and cross-patch, *α*, infection rates. For larger *β* values or larger *α* values, we found that prophylactic vaccination resulted in a smaller mean final epidemic size than when vaccines were withheld. Note that the inherent time scale of the model considered is the mean infectious period, 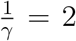 days in our examples. Therefore, the above results can be interpreted more generally by thinking of 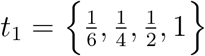 mean infectious periods as opposed to days. In this language, we have that strict-prophylactic vaccination is optimal unless the delay between infection and vaccination is less than 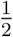 an infectious period and when the patch sizes are relatively even.

Hence, there appears to be no benefit to withholding vaccines until after infection is present in the metapopulation unless vaccination can be applied very rapidly after the first onset of infection, in which case the benefit is still minor. Instead, we find that in general for such a metapopulation with minimal structure, prophylactic vaccination results in a smaller mean final epidemic size and hence is the preferred scheme. Next, we focus on the problem of prophylactic vaccination and determining the best way of allocating a fixed number of vaccines to such a metapopulation.

## 4 Optimal prophylactic vaccination

Consider the metapopulation model described in Section 2. We know from the previous section that the preferred vaccination scheme is to allocate vaccines before the first onset of infection in the population. Hence, given a fixed number of vaccines, we consider all possible ways of allocating the available vaccines to the population and the optimal strategy is the one that results in the smallest mean final epidemic size. We first discuss how to solve this problem exactly before explaining the need for an approximation to the exact solution and considering other strategies found in the literature.

### 4.1 Optimal strategy

As the optimal strategy corresponds to the allocation with the smallest mean final epidemic size, we need to calculate the mean final epidemic size for all possible allocations of available vaccines to the population. Let *V_k_* be the number of vaccines allocated to patch *k* with the total number of vaccines, 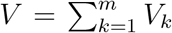, and let *u_k_* = *N_k_* – *V_k_* denote the number of unvaccinated susceptible individuals in patch *k*. Recall from Section 2 that the number of recovered (including vaccinated) individuals at the end of the epidemic for every possible initial state s, ζ(s), can be obtained simultaneously by solving,

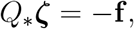

and extracting the appropriate element from ζ. Then, for the initial state of interest **u** = (*u*_1_, 0,…, *u_k_*, 0,…, *u_m_*, 0), the mean final epidemic size given the infection started in patch *k* is [ζ (**u** – **inf**_*k*_) – *V*]. By multiplying this with the probability of an individual in patch *k* becoming infected, 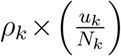, and summing over all patches, we obtain the mean final epidemic size for the initial state of interest **u**,

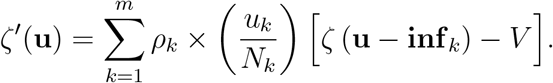

The problem of determining the optimal strategy corresponding to the minimum mean final epidemic size can then be written as the following minimisation problem,

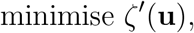

subject to

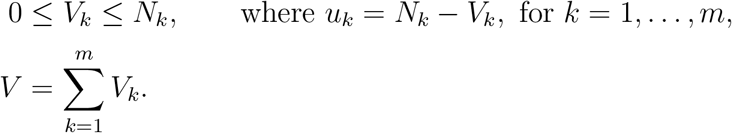

We note that this optimisation problem is relatively easy to solve once the vector ***ζ*** is calculated as it simply involves selecting the strategy corresponding to the smallest mean final epidemic size *ζ*′(**u**), which is itself a fairly trivial calculation given ***ζ***.

The difficulty with the optimisation problem lies with the calculation of the vector ***ζ*** as it is exact and requires solving a system of linear equations involving the *Q* matrix which is of order 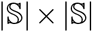. Further, the size of the state space grows with the population size as

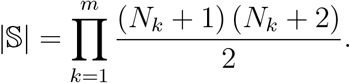

Hence, as the population size increases, obtaining the mean final epidemic size using this method becomes more computationally intensive and for large population sizes, this method is computationally intractable. This indicates a need to develop a more computationally-efficient method of estimating the mean final epidemic size to sufficient accuracy in order to develop an approximate strategy.

### 4.2 Approximate strategy

Here, we detail our approximation method which consists of two approximations. The first we call the *average initial infection rate approximation* and the second the *weakly-coupled final sizes approximation.* Also, involved in our method is a rule for determining which approximation to use for a given parameter set.

First, consider the average initial infection rate approximation. We are interested in determining the average initial infection rate for the state **u** = (*u*_1_, 0,…, *u_k_*, 0,…, *u_m_*, 0), where *u_ℓ_* = *N_ℓ_ − V_ℓ_* for *ℓ* — 1,…, *m*. The intuition is to find the vaccine allocation that minimises the average initial infection rate in order to best control the epidemic.

The average initial infection rate, *r*, for the state of interest **u**, is

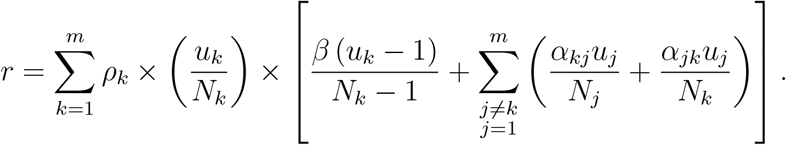

See Appendix A.2 for details of its derivation.

To determine the optimal combination of *u_k_*, for *k* = 1,…, *m*, we evaluate *r* for all feasible combinations of *u_k_* and the combination corresponding to the smallest value of *r* is chosen as the optimal.

Next, we consider the weakly-coupled final sizes approximation which is suitable when the cross-patch infection rate is sufficiently small relative to the within-patch infection rate. Recall from Section 4.1 that the problem with obtaining the optimal strategy exactly is that the calculation of the mean final epidemic size *ζ*\* (**u** − **inf**_*k*_) can be computationally intensive, or even intractable, for large population sizes. Hence, for this approximation, we develop a less computationally-intensivweakly-couplede method to approximate *ζ*\*(**u** − **inf**_*k*_).

As we assume that the cross-patch infection rate is sufficiently small relative to the within-patch infection rate, the probability of infection spreading between patches is small. Therefore, we approximate the mean final epidemic size for each patch using single population models. The basis of this approximation is to calculate the mean final epidemic size of the initially infected patch and then add to it the mean final epidemic size of each patch which has not yet experienced infection, weighted by the probability of each of those patches becoming infected. Note that we consider all possible ways (that is, different pathways of infection) the infection can spread from the initially infected patch to all fully susceptible patches.

Using this method, we obtain the approximate mean final epidemic size for each strategy. Then, the chosen strategy is the one corresponding to the smallest approximated mean final epidemic size.

As the average initial infection rate approximation and the weakly-coupled final sizes approximation each return an optimal strategy for a given parameter set, we need a method to determine which approximation, and hence corresponding strategy, is most appropriate for that parameter set. Recall that the weakly-coupled final sizes approximation is restricted to the region where the cross-patch infection rate is sufficiently small relative to the within-patch infection rate. On the other hand, there is no restriction on the average initial infection rate approximation. Hence, our method needs to determine, for a given parameter set, if the weakly-coupled final sizes approximation can be applied or the average initial infection rate approximation should be used instead.

Let us consider the transition rates of the model (described in Section 2) and focus specifically on the infection events. The rate of infection in patch *k*, for *k* = 1,…, *m*, is,

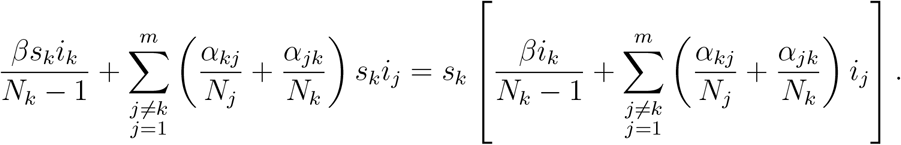

As the weakly-coupled final sizes approximation assumes that the cross-patch infection rate is sufficiently small relative to the within-patch infection rate we want, for each *k* = 1, …, *m*,

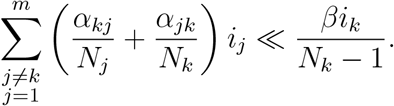

Note that the above expression is state-dependent as it depends on the values of *i_j_* and *i_k_*. Instead, we want our expression to be state-independent and so we bound these terms by the largest possible value they can take, *N_j_* and *N_k_* [14]. Thus, the expression of interest to us can be written, for each *k* = 1,…, *m*, as

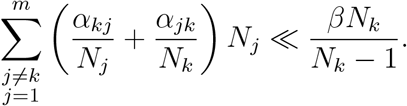

For large *N*, this expression can be approximated by,

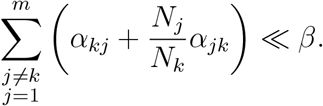

We need this condition to hold for all patches *k* = 1,…, *m*, for the weakly-coupled final sizes approximation to be a suitable approximation. Hence, for some chosen cut-off value *c* ≪ 1, if

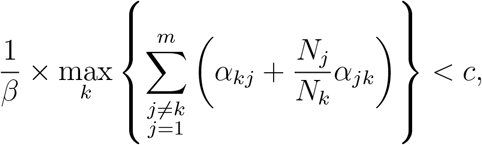

then the weakly-coupled final sizes approximation is applied. After investigation (see [14]), a cut-off value of *c* = 0.175 was chosen as a robust choice.

### 4.3 Other strategies in the literature

We compare our approximate strategy to other strategies found in the literature, in particular the *equalising* strategy [1, 2], the *deterministic* strategy, obtained from a modified model of Keeling and Shattock [3], and a *fair (pro-rata)* strategy [10], used by many as a basis for comparison.

The *equalising* strategy involves distributing vaccines in such a way that an equal, or close to equal, number of susceptible individuals remain in each household after vaccination. This was shown to be *optimal* for a stochastic SIR model with an infinitely large population of households of size two, three and four and was conjectured to hold for larger household sizes, with numerical investigations supporting this conjecture. Their definition of optimality was to determine the minimal number of vaccines and an optimal allocation scheme that brings the basic household reproduction number, *R*_0_, below 1. It was also assumed that vaccination rendered an individual immune and that the transmission of infection is density dependent, that is, the chance of transmission between any two individuals in the population is independent of the household size. Figure 3 illustrates this strategy for a simple example consisting of three households of size *N*_1_ = 2, *N*_2_ = 4, *N*_3_ = 6 with *V* = 6 vaccines.

**Figure 3:**
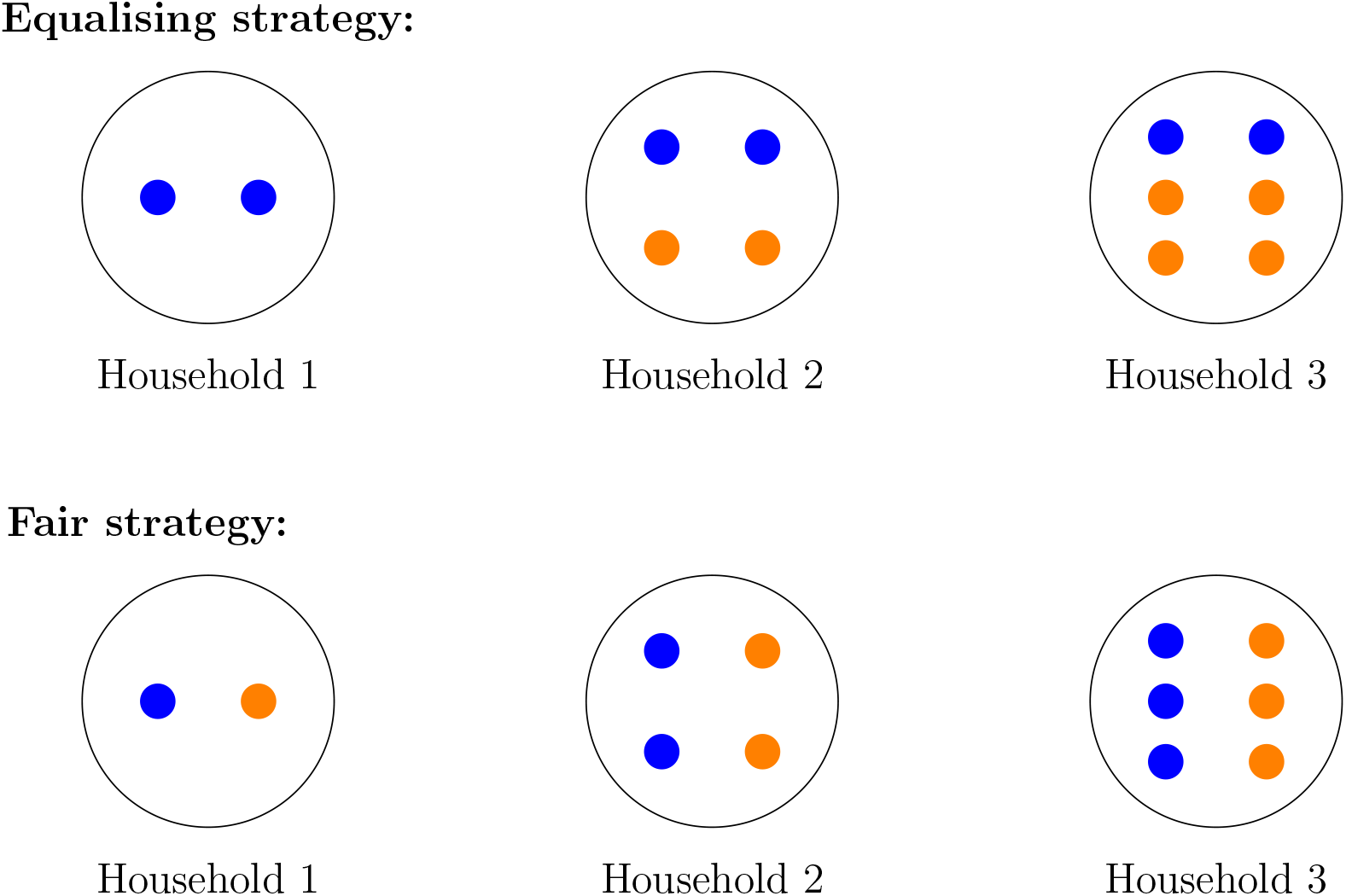
Example illustrating the allocation of vaccines using the equalising strategy and the fair strategy for a simple example with three households of size *N*_1_ = 2, *N*_2_ = 4, *N*_3_ = 6 with *V* = 6 vaccines. Here, blue represents a susceptible individual and orange represents a vaccinated individual.

Next, we consider a *fair (pro-rata)* strategy which allocates vaccines in proportion to the number of susceptible individuals in each patch. Figure 3 illustrates this strategy for a simple example consisting of three households of size *N*_1_ = 2, *N*_2_ = 4, *N*_3_ = 6 with *V* = 6 vaccines.

The final strategy we consider is the *deterministic* strategy obtained from a modified version of a deterministic model used by Keeling and Shattock [3]. For details of the modifications, see Appendix A.4. The final size equation used to determine the final size for patch *ℓ*, for *ℓ* = 1,…, *m*, where infection has been seeded in patch *k*, is,

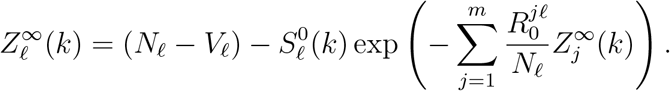

Using simple recursion, we can solve the revised final size equation by,

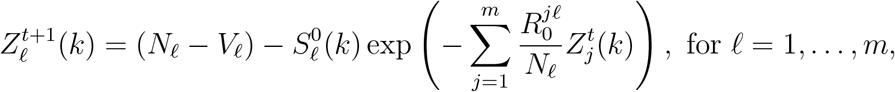

with

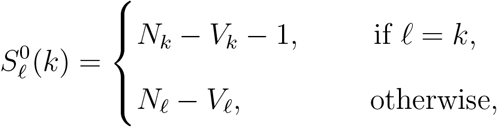

and initial condition,

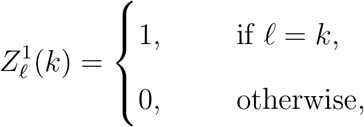

where

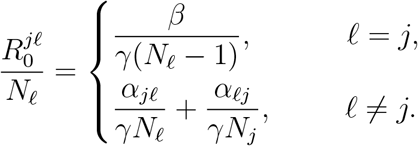

As the recursive calculation of the final size for each patch, 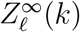, depends on the knowledge of which patch infection has been seeded in, this tells us that the final epidemic size is dependent on the initial state. Hence, for a given initial state **u**(*k*) = (*u*_1_, 0,…, *u_k_* − 1,1,…, *u_m_*, 0), where infection has been seeded in patch *k*, the final epidemic size for **u**(*k*) is,

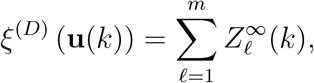

where 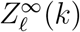 is the final size for patch *ℓ* obtained through recursion and is dependent on **u**(*k*) and hence the patch *k* where infection is seeded.

This calculation of *ξ*^(*D*)^ (**u**(*k*)) gives us the final epidemic size when infection has been seeded in patch *k*; hence, for a given parameter set, the mean final epidemic size for a given strategy **u** = (*u*_1_, 0,…, *u_k_*, 0,…, *u_m_*, 0) is,

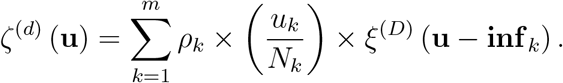

Then, the optimal deterministic strategy is the one corresponding to the smallest *ζ*^(*d*)^(**u**), calculated out of all possible strategies **u**.

### 4.4 Results

#### 4.4.1 Small populations

We first consider a small example with three patches of sizes *N*_1_ = 6, *N*_2_ = 12 and *N*_3_ = 18, with *V* = 9 vaccines available to the population. For this example, we make the assumption that the cross-patch infection rate is the same for all patches, i.e., *α_kj_* = *α* for *k,j* = 1, 2, 3 and *j* ≠ *k*. To allow for easy presentation of the results, we fix the recovery rate 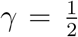 and vary the within-patch infection rate *β* ∈ [0.5, 5] and the cross-patch infection rate *α* ∈ [0.01,0.1] × *β*. Figure 4a consists of the optimal strategies and Figure 4b consists of the approximate strategies for this example. Note that as there are a large number of possible vaccine allocation schemes, we have chosen a colour map that only considers the required vaccine allocations.

**Figure 4:**
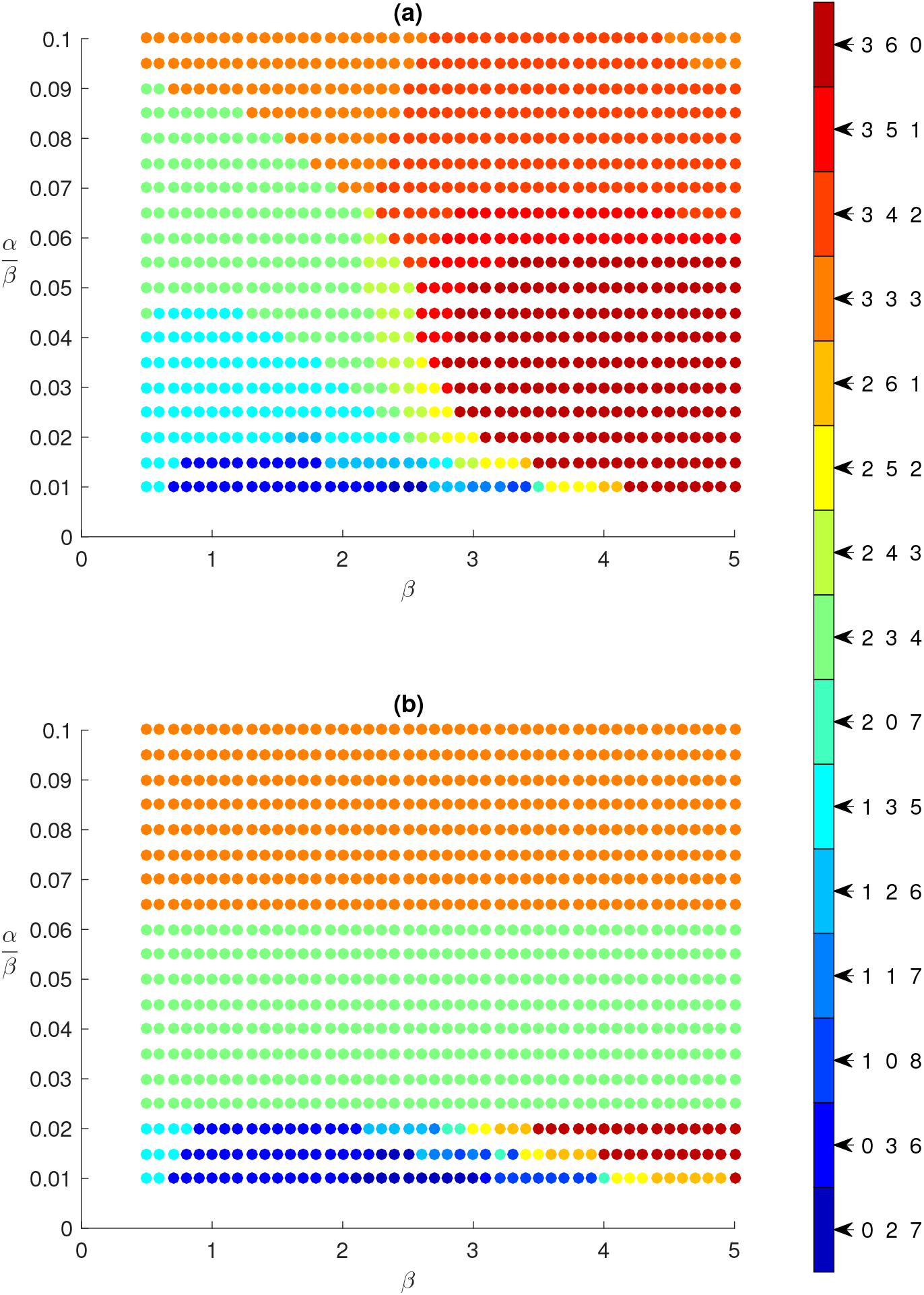
Three-patch example with *N*_1_ = 6, *N*_2_ = 12, *N*_3_ = 18, *V* = 9 vaccines, 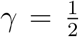, within-patch infection rate *β* ∈ [0.5, 5] and between-patch infection rate *α* ∈ [0.01,0.1] × *β*. (a): Optimal strategies, (b): Approximate strategy. The colour labels represent the number of vaccines allocated to each patch, (*V*_1_, *V*_2_, *V*_3_). Only the optimal allocations are shown in this figure. This figure shows that the optimal and approximate strategies match reasonably well especially for small 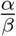 values.

We can see from this figure that there are some differences between the approximate and optimal strategies. However, we note that comparing the strategies may not be the best approach as strategies which differ by a single vaccinated individual may not result in a large difference in the mean final epidemic size. Instead, we use the relative difference in mean final epidemic size as a measure to determine how well a proposed strategy performs in comparison with the optimal strategy.

Let *ζ_opt_* denote the mean final epidemic size for the optimal strategy and *ζ_prop_* denote the mean final epidemic size for the proposed strategy. Note that the mean final epidemic sizes, *ζ_opt_* and *ζ_prop_*, are calculated exactly using the method in Section 4.1. Then, the relative difference is given by,

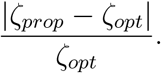

Note that this is the relative difference for a single parameter set. If we have a range of parameters and are interested in comparing between various proposed strategies, then a single number which describes how well a proposed strategy performs over the range of parameters is preferred. Hence, we have chosen to use the average and maximum relative differences as our measures for comparison between proposed strategies.

Figure 5 consists of the relative difference between each proposed strategy (approximate, deterministic, equalising and fair) and the optimal strategy and Table 1 consists of the average and maximum relative differences. Note that in Table 1, we consider two parameter regions. The first region consists of the entire parameter range and the second region has the additional condition of *β* ∈ [0.5, 0.8] which corresponds to influenza-like parameters, i.e., *R*_0_ E [1,1.6].

**Figure 5:**
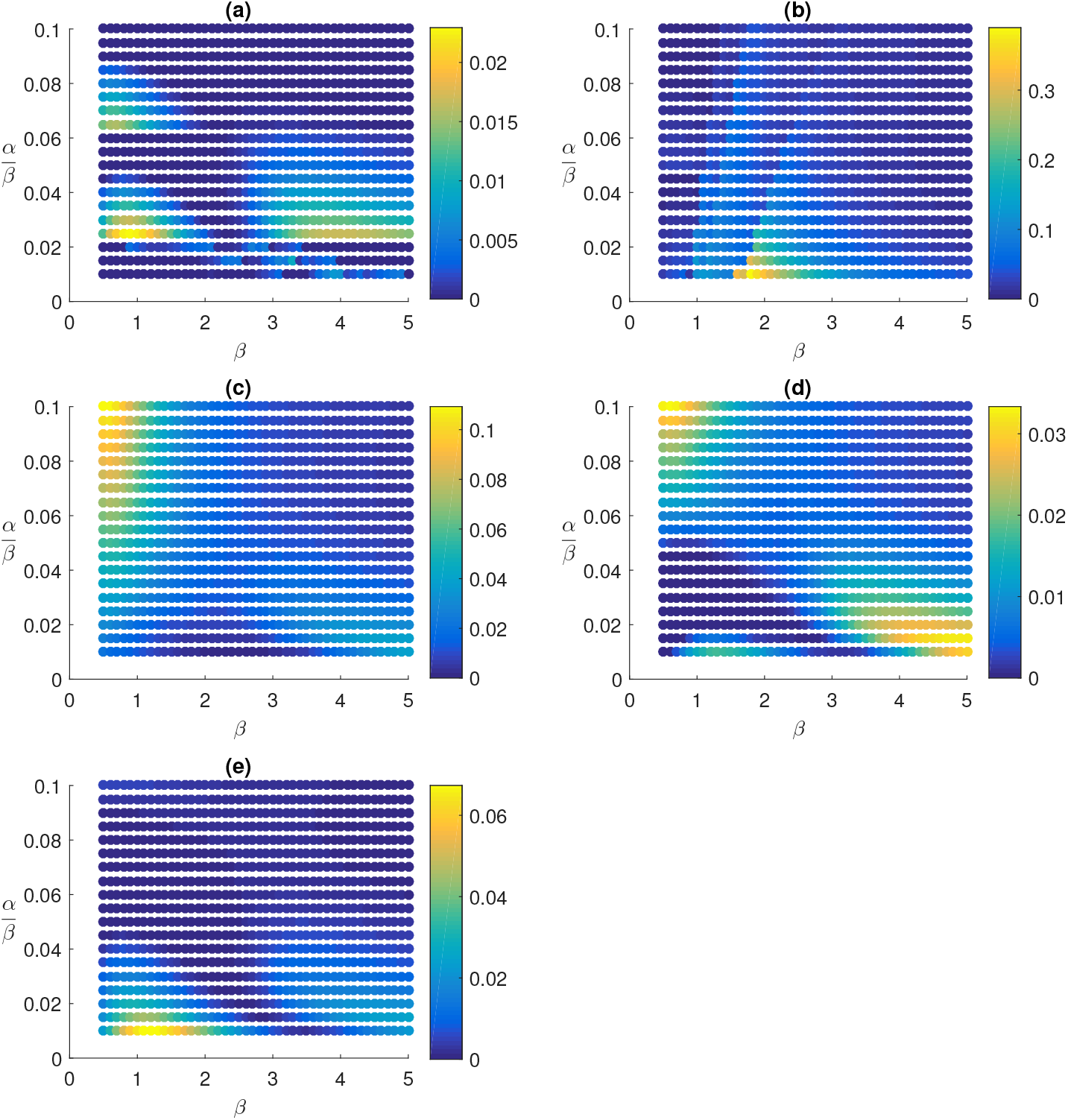
Relative difference in mean final epidemic size between the different strategies and the optimal for the three-patch example with *N*_1_ = 6, *N*_2_ = 12, *N*_3_ = 18, *V* = 9 vaccines, 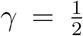, within-patch infection rate *β* ∈ [0.5, 5] and between-patch infection rate *α* ∈ [0.01,0.1] × *β*. (a): Approximate strategy versus optimal, (b): Deterministic strategy versus optimal, (c): Equalising strategy (*V*_1_, *V*_2_, *V*_3_) = (0, 2, 7) versus optimal, (d): Fair strategy (*V*_1_, *V*_2_, *V*_3_) = (1, 3, 5) versus optimal, (e): Fair strategy (2, 3, 4) versus optimal. Note each subfigure has its own colour scale. Further, note that two fair strategies are possible due to the rounding of proportions/individuals. This shows that the approximate strategy compares well with the optimal strategy and performs better than the other strategies considered.

We observe from both Figure 5 and Table 1 that the approximate strategy performs considerably better than the deterministic and equalising strategies and is also slightly better than the fair strategy for both parameter regions considered. Further, we also note that the average and maximum relative differences for the approximate strategy are very small in magnitude. Hence, this gives us confidence that the approximate strategy performs very well when compared to other strategies.

Another important point to note is the amount of time required to obtain the optimal strategies. Both the exact method and the approximate method were run on a Lenovo NeXtScale system consisting of 120 nodes, where one core of an Intel^®^ 2.3 GHz Xeon E5-2698v3 node with 12 GB of RAM was used. The exact method required 3313 seconds to obtain the optimal strategies whilst the approximate method took only 1.74 seconds to obtain the approximate strategies. This considerable speed up in computational time combined with the small relative differences enhances the attractiveness of the approximate method. Further, as the size of the patches increases, obtaining the optimal strategies exactly becomes computationally infeasible, making an approximation method necessary to determine the best strategies.

**Table 1:** Average and maximum relative difference for each proposed strategy for the three-patch example with *N*_1_ = 6, *N*_2_ = 12, *N*_3_ = 18, *V* = 9 vaccines and 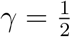.

| Proposed<br>strategy | Region | Relative Difference |  |
| --- | --- | --- | --- |
|  |  | Average | Maximum |
| Approximate | Full | **0.0027** | **0.0229** |
| Deterministic | Full | 0.0363 | 0.3895 |
| Equalising | Full | 0.0212 | 0.1093 |
| Fair (1,3,5) | Full | 0.0074 | 0.0333 |
| Fair (2,3,4) | Full | 0.0058 | 0.0673 |
| Approximate | Influenza-like | **0.0047** | **0.0222** |
| Deterministic | Influenza-like | 0.0127 | 0.0509 |
| Equalising | Influenza-like | 0.0610 | 0.1093 |
| Fair (1,3,5) | Influenza-like | 0.0098 | 0.0333 |
| Fair (2,3,4) | Influenza-like | 0.0074 | 0.0509 |

#### 4.4.2 Larger populations

Next, we explore an example of a larger problem where it is computationally infeasible to obtain the optimal strategies exactly. Consider an example with 1800 individuals split into three patches of sizes *N*_1_ = 300, *N*_2_ = 600 and *N*_3_ = 900, and *V* = 450 vaccines. Note that this is exactly the example used in the previous section, but scaled up by a factor of 50. As with the previous example, we fix the recovery rate for all patches 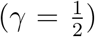 and assume that the within-patch infection rate is the same for all patches and the cross-patch infection rate is also the same for all patches. We also choose to vary the within-patch infection rate *β* ∈ [0.5, 5] and the cross-patch infection rate, *α* ∈ [0.01, 0.1] × *β*. Figures 6a and 6b consist of the approximate and deterministic strategies for each (*α, β*) pair respectively. From these figures, we observe that the approximate strategy differs from the deterministic strategy for *β* > 1.5.

**Figure 6:**
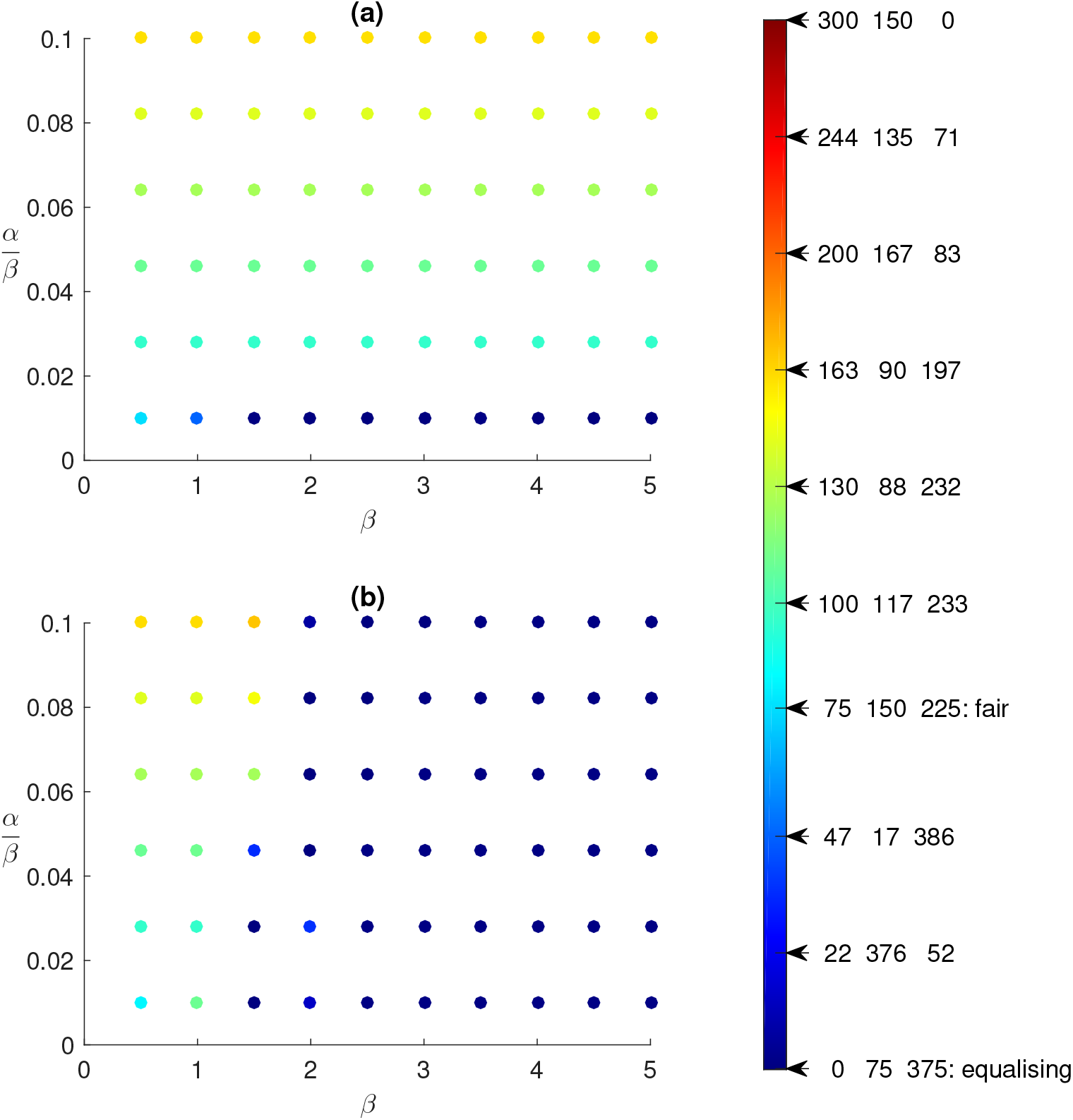
Optimal strategies for the large three-patch example with *N*_1_ = 300, *N*_2_ = 600, *N*_3_ = 900, *V* = 450 vaccines, 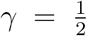, within-patch infection rate *β* ∈ [0.5, 5] and between-patch infection rate *α* ∈ [0.01,0.1] × *β*. (a): Approximate strategy, (b): Deterministic strategy. The colour labels represent the number of vaccines allocated to each patch, (*V*_1_, *V*_2_, *V*_3_). This figure shows that the approximate and optimal strategies differ for *β* > 1.5.

Unlike the previous example, it is computationally infeasible to calculate the mean final epidemic size exactly for this problem. Hence, we are unable to use the previous method of comparison where we calculated the relative difference in mean final epidemic size between the optimal strategy and the proposed strategies (approximate, deterministic, equalising and fair). Instead, we perform statistical tests to determine if the mean final epidemic size between the strategies are significantly different. This can then tell us how well the approximate strategy performs in comparison to the deterministic, equalising and fair strategies. We highlight some key points of our method of comparison and refer the reader to Appendix A.5 for more details.

As it is computationally infeasible to calculate the mean final epidemic sizes for this example exactly, we use Sellke’s simulation method [19] to estimate the final epidemic size for a given (*α, β*) pair and a given strategy (initial state).

The first statistical test we perform is the one-way ANOVA test, which tests if the means for all strategies are the same. For our problem, we have chosen to perform the one-way ANOVA test at a 5% significance level using the `aov` function in R [20]. If we fail to reject the null hypothesis for the one-way ANOVA test, then we conclude that there is insufficient evidence that the mean final epidemic sizes of the strategies are significantly different. Otherwise, the null hypothesis is rejected and we perform pairwise comparisons using Dunnett’s test [21, 22], at the 5% significance level with the `glht` function in R [23], to determine which strategy or strategies differ from the approximate strategy. If the null hypothesis for Dunnett’s test is rejected, then we conclude that there is a difference in the mean final epidemic size between the approximate strategy and the other strategy of interest. Further, if the difference (other – approximate) in the mean final epidemic sizes is positive, then the approximate strategy is better than the other strategy. Otherwise, the other strategy is better.

We perform this testing procedure for the (*α, β*) pairs considered, each with 10^6^ simulations. Figure 7 displays the results of the one-way ANOVA test. We observe from this figure that for all (*α, β*) pairs, at least one of the mean final epidemic sizes is significantly different from the other means at the 5% significance level. Hence, we perform pairwise comparisons, using Dunnett’s test, to determine whether the approximate strategy performs better than the other strategies. Figure 8 consists of the difference in sample means between the deterministic, equalising and fair strategies with the approximate strategy, as well as the p-values from Dunnett’s test for each pairwise comparison. By showing both the difference in sample means as well as the p-values, we can use these figures to determine regions where the means are significantly different (for any choice of significance level) and also determine if the other strategies outperform the approximate strategy. Figure 9 combines the results of Dunnett’s test at a 5% significance level with the difference in sample means for each pairwise comparison. It indicates where the other strategies are significantly different from the approximate strategy as well as how different the sample means of the strategies are through the intensity of the colour. Figure 10 combines the results of the individual Dunnett’s test at a 5% significance level into a single figure. From these figures, we observe that the approximate strategy performs well when compared to the equalising and fair strategies.

**Figure 7:**
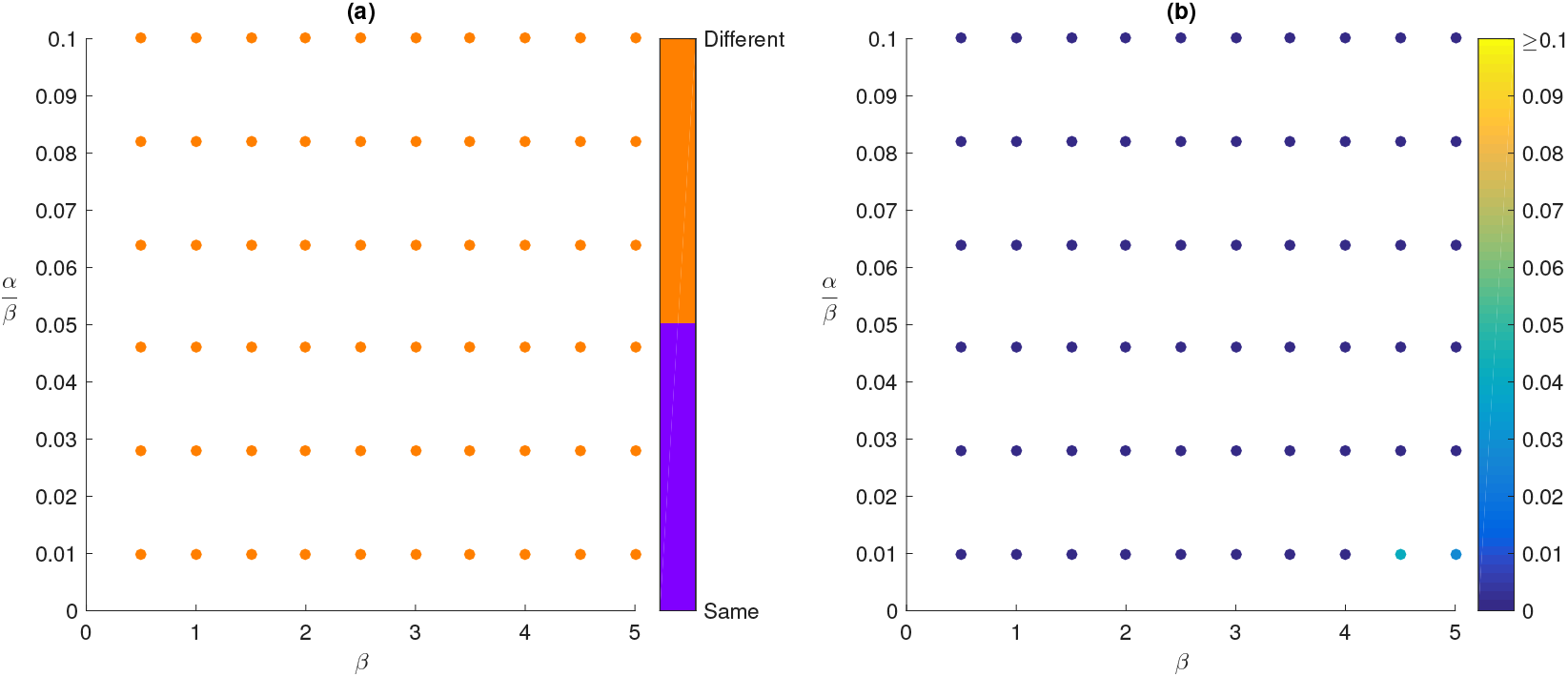
Comparison to determine if the means of all strategies are the same using a one-way ANOVA test for the large three-patch example with *N*_1_ = 300, *N*_2_ = 600, *N*_3_ = 900, *V* = 450 vaccines and 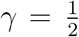. (a): Indicates whether all means are the same or at least one mean is different at a 5% significance level, (b): p-values for the one-way ANOVA test.

**Figure 8:**
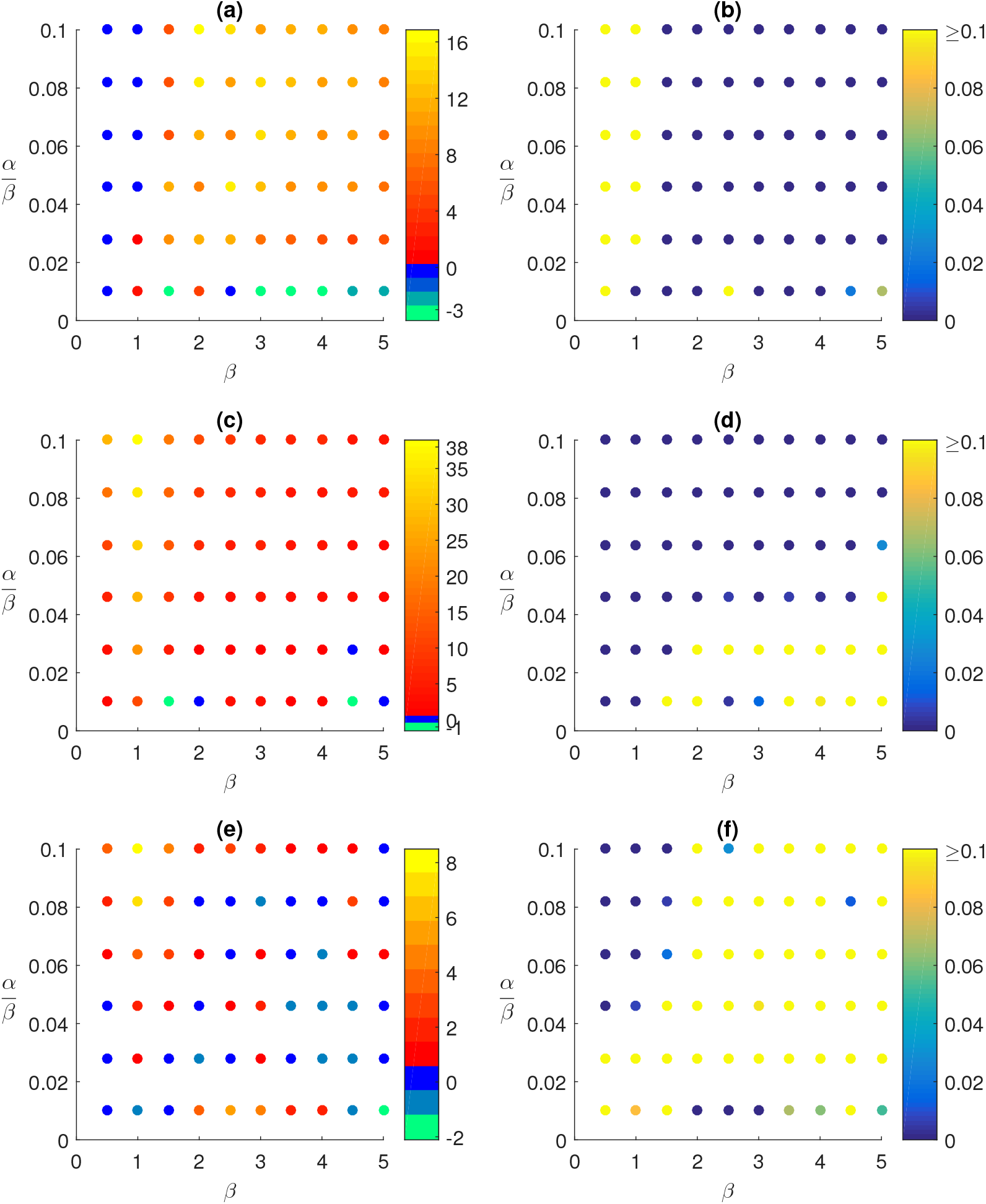
Comparison to determine whether the other strategies are better than the approximate strategy using Dunnett’s test for the large three-patch example with *N*_1_ = 300, *N*_2_ = 600, *N*_3_ = 900, *V* = 450 vaccines and 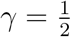. (a): Difference in sample means (deterministic – approximate), (b): p-values for Dunnett’s test, (c): Difference in sample means (equalising – approximate), (d): p-values for Dunnett’s test, (e): Difference in sample means (fair – approximate), (f): p-values for Dunnett’s test.

When comparing with the deterministic strategy, we observe from the figures that the deterministic strategy appears to perform better than the approximate strategy in the region where 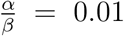. However, when we consider Figure 9a, which combines the results of Dunnett’s test at a 5% significance level with the difference in sample means, we see that when the approximate strategy performs better than the deterministic strategy, the difference in sample means is larger than the difference when the deterministic strategy performs better. Hence, over the range of (*α, β*) pairs considered, it appears that the approximate strategy is more robust than the other strategies.

**Figure 9:**
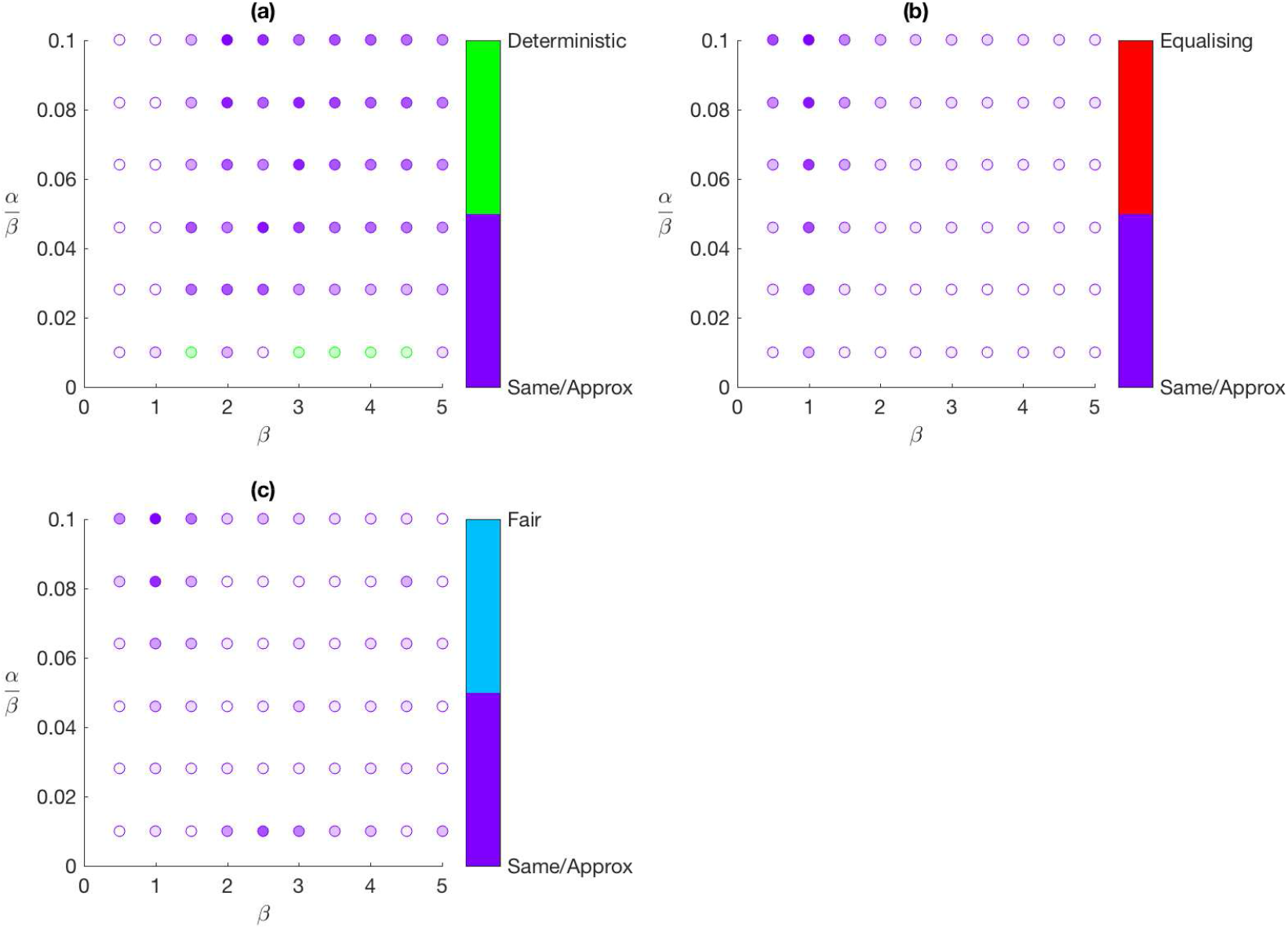
Comparison of the approximate strategy with other strategies using Dunnett’s test at a 5% significance level for the large three-patch example with *N*_1_ = 300, *N*_2_ = 600, *N*_3_ = 900, *V* = 450 vaccines and 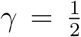. (a): Indicates the deterministic strategy is better (green) or the approximate strategy is better or there is no difference between strategies (purple), (b): Indicates the equalising strategy is better (red) or the approximate strategy is better or there is no difference between strategies (purple), (c): Indicates the fair strategy is better (blue) or the approximate strategy is better or there is no difference between strategies (purple). The intensity of the colour increases with the difference in sample means.

**Figure 10:**
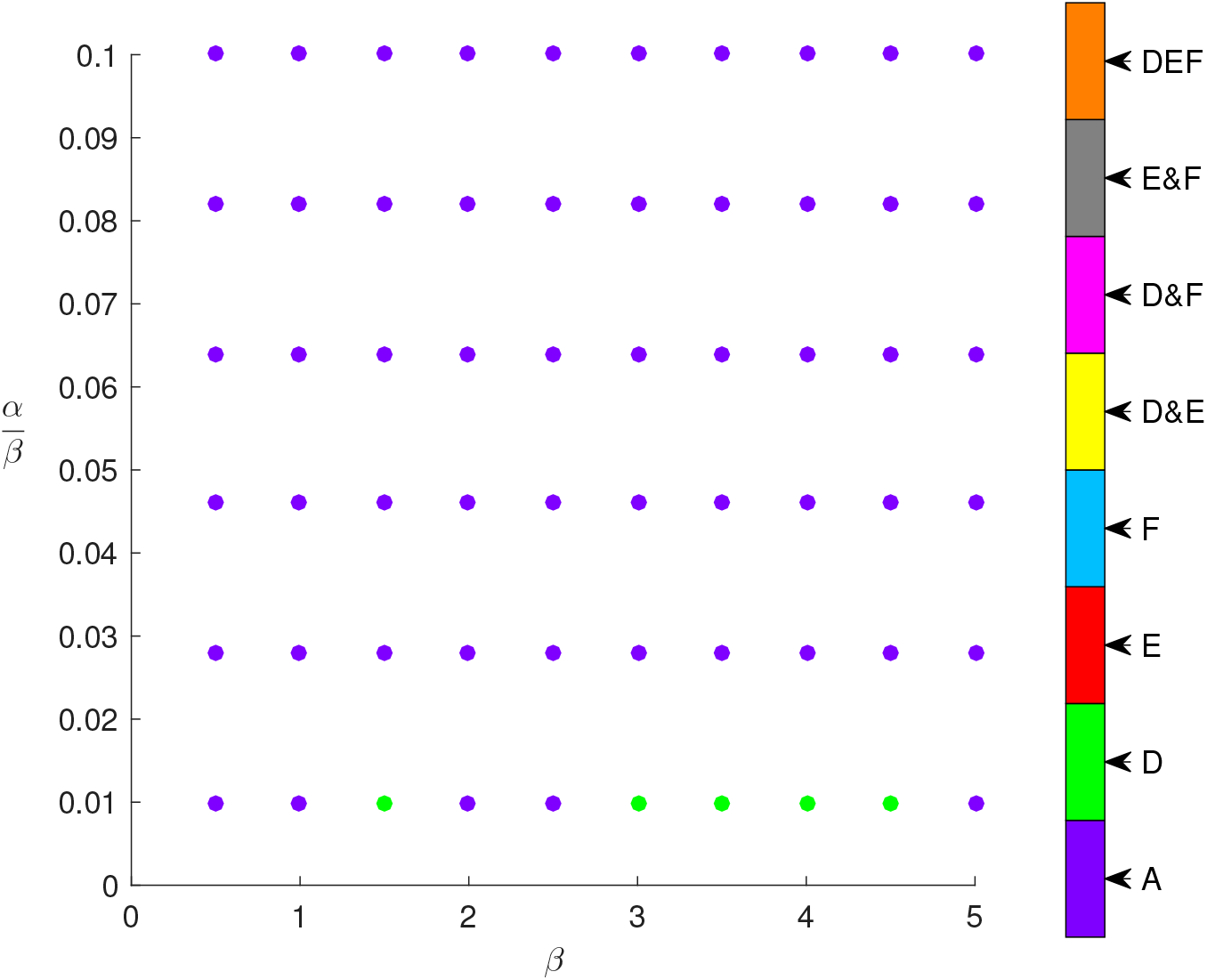
Combines individual tests between the approximate strategy and other strategies using Dunnett’s test at a 5% significance level for the large three-patch example with *N*_1_ = 300, *N*_2_ = 600, *N*_3_ — 900, *V* = 450 vaccines and 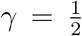. The legend indicates which strategies are better than the approximate strategy, where A, D, E and F represent the approximate strategy, the deterministic strategy, the equalising strategy and the fair strategy, respectively. Note that purple indicates either the approximate strategy performed better or there was no difference between the performance of the approximate strategy and that of the other strategies.

## 5 Discussion

The question of how best to allocate a limited supply of vaccines to a population to minimise the expected number of people that become infected over the course of an epidemic is of great importance to public health officials. We addressed this question in two parts, when to allocate and where to allocate. For both parts, we utilised a stochastic SIR metapopulation model and assumed that the entire population is initially susceptible with infection occurring through a single attempted import into the population.

We first explored the question of when best to allocate vaccines, before infection or after infection is present in the population, and considered three different allocation strategies: strict-prophylactic vaccination, predominantly-prophylactic vaccination and reactive vaccination. Through the use of a modified backward dynamic programming algorithm which included a delay between the first onset of infection and vaccination, we found that in general, strict-prophylactic vaccination resulted in a smaller mean final epidemic size. We believe this to hold for any metapopulation which lacks significant structure which might be exploited. Part of the reason for the success of the prophylactic scheme here is its accounting for import of infection in the objective for determining the allocation. Hence, for the next question of where best to allocate a limited supply of vaccines to a metapopulation, we focused on the case of strict-prophylactic vaccination.

Determining the optimal strategy for a given set of parameters can be a computationally intensive task especially as the population size increases; for large population sizes, this becomes computationally intractable. Hence, we developed a more computationally-efficient method to obtain an approximately optimal strategy. Our approximation method consists of two approximations, the average initial infection rate approximation and the weakly-coupled final sizes approximation, as well as a rule for determining which approximation to use for a given parameter set. We considered two examples, a small population size and a larger population size, and compared the performance of our approximate strategy with other strategies in the literature (deterministic, equalising and fair). In general, we observed that our approximate strategy performed well when compared to the other strategies and also appeared to be more robust than the other strategies for a range of parameters.

These examples provide us with confidence that the approximate strategy performs well across examples of different sizes. Overall, we note that whenever the optimal strategy can be obtained, it should, of course, be used. However, as the population size increases, we are unable to obtain the optimal strategy and so the approximate strategy we developed should be used instead. Then, if we consider even larger population sizes, there appears to be some benefit in using the deterministic strategy.

This work can be extended in a number of ways. One possible extension is to consider other types of vaccination. Here, we have assumed that vaccination renders an individual immune to the disease. However, this may not be entirely realistic. Some vaccines may not be completely effective and so only a proportion of vaccinated individuals are immune to the disease (*all-or-nothing* vaccination), or perhaps, vaccination might only reduce an individual’s chance of becoming infected (*leaky* vaccine). To incorporate all-or-nothing vaccination in our model, binomial probabilities would need to be included and all possible combinations of each strategy considered. While for leaky vaccines, the size of the state space of our model would increase to account for the additional class of susceptible but vaccinated individuals. Both of these variations to vaccination can be incorporated relatively easily in our model. However, they increase the computational effort required to determine the optimal solution and so, other simplifications and approximations may be needed for useful results to be obtained.

Another possible extension is to obtain strategies which are robust to different objectives (such as the duration of an epidemic or the peak (maximum number of infected individuals) of an epidemic). This could be explored by performing multicriteria optimisation and weighting multiple objectives. The allocation of other resources, such as anti-virals or surveillance resources, could also be investigated in a similar way.

## A Appendix

### A.1 Modified BDP algorithm

The modified BDP algorithm utilises backward dynamic programming [17, 18] to determine an optimal vaccination strategy. Here, we discuss the terms in the backward dynamic programming algorithm and how they relate to our problem. Note that we have chosen to consider the allocation of vaccines as a finite-horizon problem where after a given time *T*, no more vaccines are allocated and the process evolves without any intervention. We let *F_t_*(**S**_*t*_, *v*) represent the minimum mean final epidemic size starting in state **S**_*t*_ at time *t* with optimal actions chosen from time *t* until the horizon *T* and with *v* vaccines available. At the time-horizon *T*, we know that *F_T_*(**S**_*T*_, *v*) is the mean final epidemic size for state **S**_*T*_ as no actions are taken from the horizon *T*.

As highlighted in Section 3, we have chosen to simplify the modelling and limit the number of possible actions taken at each time step by only allowing at most one vaccine to be allocated at each time step during the infection. That is, the possible actions at time step *t* are:

- *a_t_* = 0 = no vaccine allocated,
- *a_t_* = *k* = a single vaccine allocated to patch *k*, for *k* = 1,…, *m*.

Note that if a vaccine is not allocated, then it is stockpiled for use at a later time step. Hence, for a given state **S**_*t*_, the set of possible actions are given by, 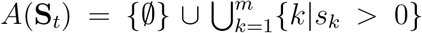, as there needs to be at least one susceptible individual in patch k for vaccination to occur in that patch. After an action is taken, the process evolves to the next time step according to *P*(s′|**S**_*t*_, *a_t_*), which is the probability of the process moving to state s′, given the current state **S**_*t*_ and action *a_t_* is chosen. This probability can be obtained from the transition probability matrix, *P*(*t*′) = *e^Qt′^*, which consists of the probabilities of moving from one state to another in time *t*′, for all *t*′ ≥ 0.

Recall from Section 3 that our simplification of only allowing at most one vaccine to be allocated at each time step does not restrict the model to only allocating at most one vaccine each day as a time step, *t*′, does not necessarily correspond to a day. If we assume that a day corresponds to a time unit of 1, then smaller values of *t*′ result in the process evolving over smaller time steps (than a day) and so vaccines are allocated at a faster rate. For our example, we let 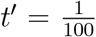 which essentially means that vaccines are able to be allocated almost simultaneously. In Algorithm 1, 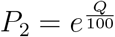 is the transition probability matrix governing the evolution of the process after the attempted import of infection and when vaccination is allowed to be applied.

The other term we detail is C(**S**_*t*_, *a_t_*), which is the contribution of choosing action *a_t_*, given the process is in state **S**_*t*_ at time *t*. If no vaccine is allocated, that is, action *a_t_* = 0 is chosen, then *C*(**S**_*t*_, 0) = 0, for all *t* and 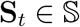, as there is no change in the mean final epidemic size. On the other hand, for all other actions *a_t_* = *k* for *k* = 1,…, *m*, an individual in patch k is vaccinated and moved to the recovered class. As the method for obtaining the mean final epidemic size in the backward dynamic programming algorithm determines the number of recovered individuals, we need to remove the vaccinated individual from the mean final epidemic size calculation and so *C*(**S**_*t*_, *a_t_*) = −1, for all *t*, 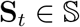 and *a_t_* ∈ *A*(**S**_*t*_)\{0}.

Next, we describe the post-processing step to account for prophylactic vaccination and the attempted import of infection. We note here that unlike vaccination after infection is present in the population, vaccines allocated prophylactically are distributed simultaneously before the attempted import of infection as opposed to a single vaccine each time step. To allow for prophylactic vaccination, we assume that at the initial time *t* = 1, the entire population is susceptible. Hence, the initial state **S**_1_ = (*N*_1_, 0, …, *N_k_*, 0,…, *N_m_*, 0). We also assume that the attempted import of infection occurs at time *t* = 1 and any number of vaccines, up to the maximum available *V*, can be allocated to the population before infection. Then, to determine the optimal allocation of these vaccines before infection, we consider all possible ways of allocating the vaccines to the patches as well as the option of withholding the vaccines to be allocated post-infection. We use *A_pre_*(**S**_1_, *V*) to denote this set of possible actions for state **S**_1_ and vaccines *V*. To better understand *A_pre_*(**S**_1_, *V*), consider a simple example with two patches, initial state **S**_1_ and *V* = 2 vaccines. For this example, the set *A_pre_*(**S**_1_, *V*) consists of the following actions,

- no vaccines pre-allocated and two vaccines available post infection,
- one vaccine pre-allocated to patch 1 or patch 2 and one vaccine available post infection,
- one vaccine pre-allocated to each patch and no vaccines available post infection, and
- two vaccines pre-allocated to patch 1 or patch 2 and no vaccines available post infection.

Note that this set comprises of all three cases, strict-prophylactic vaccination, predominantly-prophylactic vaccination and reactive vaccination, as well as all possible ways of allocating vaccines within each case. Then, to determine the best vaccination scheme, we compare the mean final epidemic sizes of all possible schemes in the set *A_pre_*(**S**_1_, *V*).

The general backward dynamic programming algorithm returns the optimal mean final epidemic size for each state in the state space without accounting for the attempted import of infection or prophylactic vaccination. Let *F*_1_(**S**, *v*) be the minimum mean final epidemic size starting in state 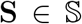 at time *t* = 1 with v vaccines available, for *v* = 0,…, *V*, obtained from the general backward dynamic programming algorithm. Then, let 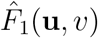 be the minimum mean final epidemic size of starting in state **u** = (*u*_1_, 0,…, *u_k_*, 0,…, *u_m_*, 0), at time *t* = 1 with *v* vaccines, for *v* = 0, …, *V*, after accounting for the attempted import of infection in the population and prophylactic vaccination. Note that *u_k_* = *N_k_* = *V_k_* is the number of unvaccinated susceptible individuals in patch *k* and *V_k_* is the number of vaccines allocated to patch *k* with 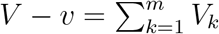. Thus, we define,

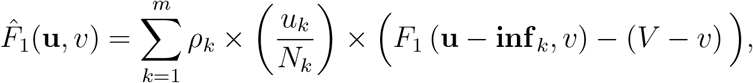

for *v* = 0,…, *V*, where **inf**_*k*_ = **e**_2*k*_ = **e**_2*k*−1_ and **e**_*k*_ is a vector of 0s with a 1 in the *k*^th^ position. As the method used to determine the mean final epidemic size for the backward dynamic programming algorithm contains vaccinated individuals, we subtract the number of pre-allocated vaccines (*V* − *v*) from the mean final epidemic size.

We can determine the minimum mean final epidemic size under the optimal vaccination scheme, for the initial state **S**_1_ = (*N*_1_, 0,…, *N_k_*, 0,…, *N_m_*, 0) and a given number of available vaccines *V*, in the following way,

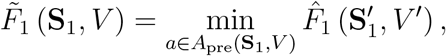

where 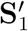 is the state of the process and *V*′ is the number of vaccines remaining after allocation *a* ∈ *A*_pre_ (**S**_1_, *V*) has been applied. The optimal vaccination scheme is the allocation a that solves the above minimisation problem.

Algorithm 1 details the modified BDP algorithm used to determine the optimal vaccination strategy. Note that in Algorithm 1, *t*′ represents the chosen time step for the problem and *P*_1_ = *e*^*Qt*_1_^ is the transition probability matrix containing the probabilities of moving from one state to another in time *t*_1_, where *t*_1_ is the delay between the first onset of infection in the population and non-prophylactic vaccination.

### A.2 Average initial infection rate

Recall, the probability of a randomly chosen individual in patch *k* becoming infected is 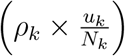, where 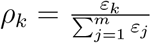, and if infection is successful, the process moves to the state (*u*_1_, 0,…, *u_k_* − 1, 1,…, *u_m_*, 0). From this state, the process can move to m possible states of further infection (ignoring the state corresponding to recovery) according to the rates specified in Section 2. Explicitly, the state (*u*_1_, 0,…, *u_k_* − 1, 1,…, *u_m_*, 0) can move to the following states with the corresponding rates,

- (*u*_1_, 0,…, *u_j_*, 0, *u_k_* = 2, 2,…, *u_m_*, 0) at rate

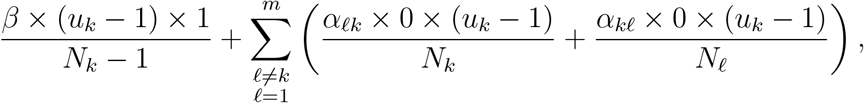
- (*u*_1_, 0,…, *u_j_* = 1, 1, *u_k_* = 1, 1, …, *u_m_*, 0) at rate

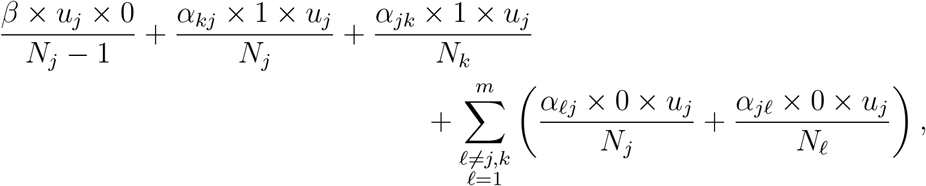

for *j* = 1,…, *m* and *j* ≠ *k*.

Hence, the initial infection rate for patch *k* is,

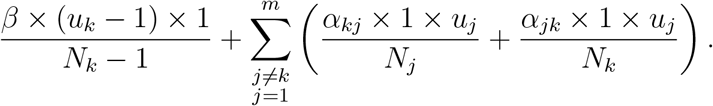

#### Algorithm 1

Backward dynamic programming algorithm modified to allow for prophylactic vaccination, attempted import of infection, choice in the rate of vaccination post infection and delay between infection and vaccination.

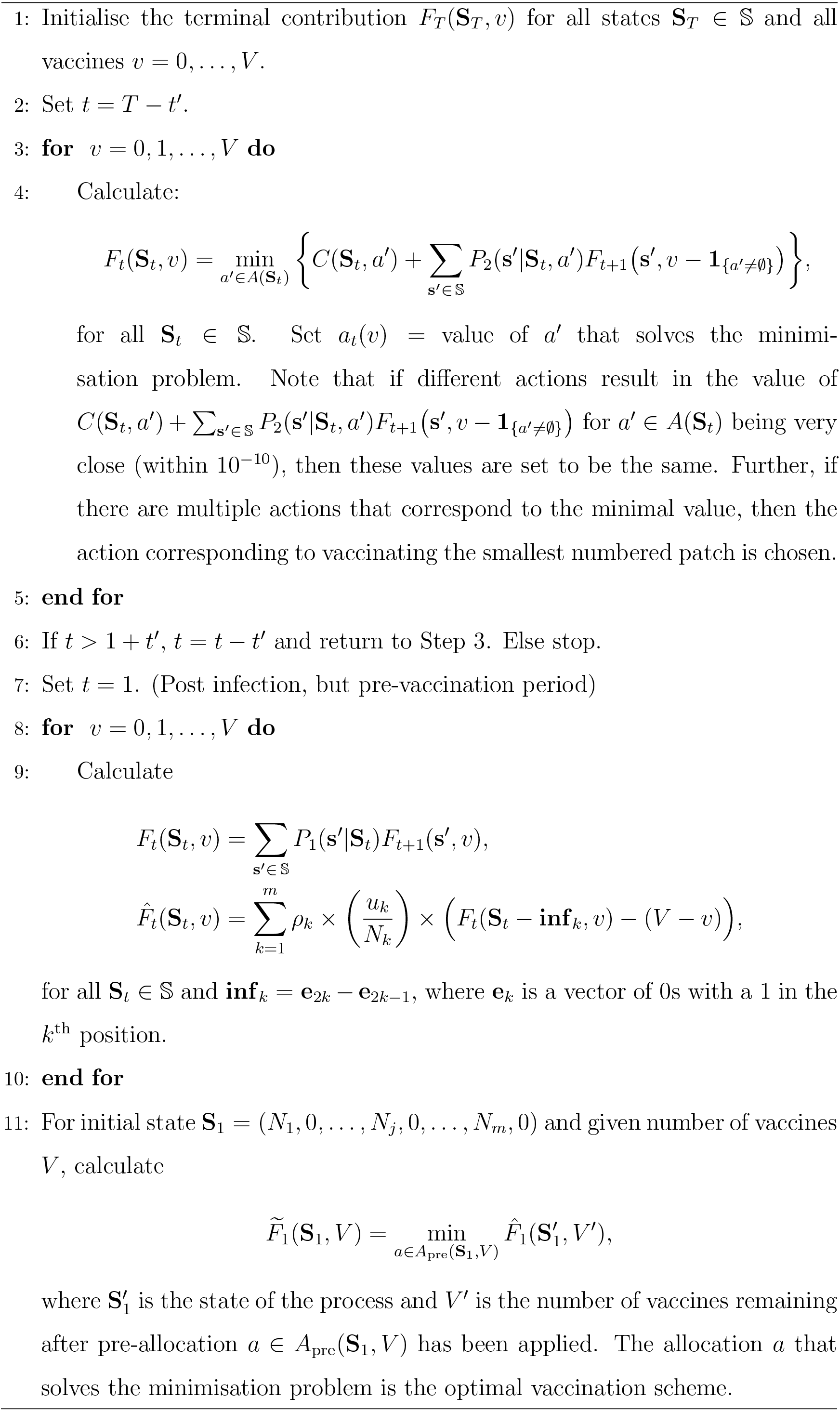

Then, to obtain the average initial infection rate, *r*, for the state of interest **u** = (*u*_1_, 0, …, *u_k_*, 0,…, *u_m_*, 0), we combine all *m* possible cases of initial infection and obtain,

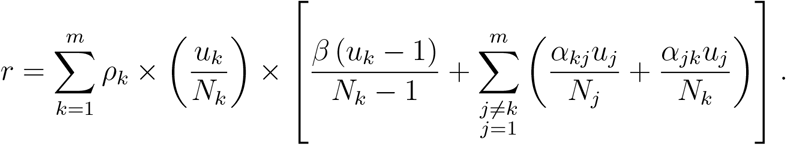

### A.3 Weakly-coupled final sizes approximation

The weakly-coupled final sizes approximation is suitable when the cross-patch infection rate is sufficiently small relative to the within-patch infection rate. Recall from Section 4.1 that the mean final epidemic size for the initial state **u** = (*u*_1_, 0, …, *u_k_*, 0, …, *u_m_*, 0) can be calculated as follows,

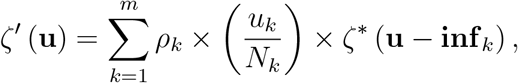

where *ζ*\* (**u** − **inf**_*k*_) = *ζ*(**u** − **inf**_*k*_) − *V*. The problem with obtaining the optimal strategy exactly, as identified previously, is that the calculation of the mean final epidemic size *ζ*\* (**u** − **inf**_*k*_) can be computationally intensive, or even intractable, for large population sizes. For this approximation, we develop a less computationally-intensive method to approximate *ζ*\* (**u** − **inf**_*k*_).

Note that we have mentioned that this approximation is appropriate when the cross-patch infection rate is sufficiently small relative to the within-patch infection rate. Under this condition, the probability of infection spreading between patches is small. Hence, we can approximate the mean final epidemic size for each patch individually, using single population models, and then combine these in an appropriate way to obtain the mean final epidemic size for a given initial state. This reduces the computational complexity of our model, as there is a significant reduction in the state space when using single population models.

Let *ζ*\* (*s_k_, i_k_*) denote the mean final epidemic size, excluding vaccinated individuals, for a single population of size *N_k_* with *s_k_* susceptible individuals and i_k_ infectious individuals. Recall that the entire population is initially susceptible and a randomly chosen individual becomes infected. Assuming that the infected individual belongs to patch *k*, the mean final epidemic size for patch *k*, obtained from a single population model, is *ζ*\* (*u_k_* − 1, 1). We then consider each *fully* susceptible patch *j* and add its mean final epidemic size, weighted by the probability of patch *j* becoming infected, to the mean final epidemic size for patch *k*. Note that we consider all possible ways (that is, different pathways of infection) the infection can spread from the initially infected patch to all fully susceptible patches. As we have assumed that the cross-patch infection rate is sufficiently small relative to the within-patch infection rate it is unlikely for previously infected patches to be re-infected. Hence, we do not consider re-infection into patches that have already been infected. Further details can be found in [14].

Next, we briefly detail the calculation of the probability of a fully susceptible patch becoming infected. Assume that patch *k* is the currently infected patch and patch *j* is a fully susceptible patch. Let *T_I_* denote the set of previously infected patches, excluding patch *k*, and let *T_S_* denote the set of fully susceptible patches. Also, let (*k* → *j* | *T_I_, T_S_*) denote the infection event from patch *k* to patch *j* ∈ *T_S_*, given the previously infected patches in *T_I_* and the fully susceptible patches in *T_S_*. Then, we have,

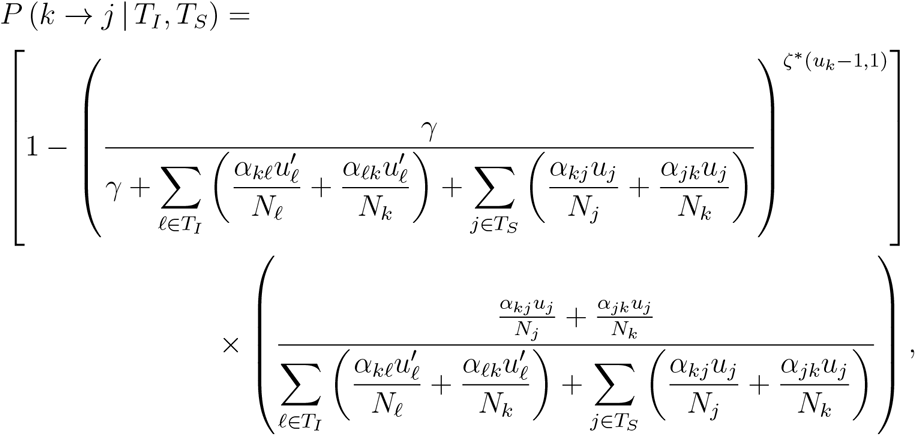

where 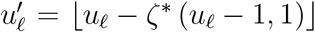, since we require the number of susceptible individuals to be an integer value. This probability is the product of the probability that at least one infectious individual in patch *k* infects a susceptible individual in another patch and the conditional probability of one infectious individual in patch *k* infecting patch *j* ∈ *T_s_* instead any other patch, given that cross-infection occurs.

Here, we provide a detailed example of the approximation for *m* = 3 patches. For this case, the quantities of interest to us are *ζ*\* (*u*_1_ − 1, 1, *u*_2_, 0, *u*_3_, 0), *ζ*\* (*u*_1_, 0, *u*_2_ − 1, 1, *u*_3_, 0) and *ζ*\* (*u*_1_, 0, *u*_2_, 0, *u*_3_ − 1, 1). We consider *ζ*\* (*u*_1_ − 1, 1, *u*_2_, 0, *u*_3_, 0) and note that the other two expressions are analogous, *ζ*\* (*u*_1_ = 1, 1, *u*_2_, 0, *u*_3_, 0) = *ζ*\* (*u*_1_ = 1, 1)

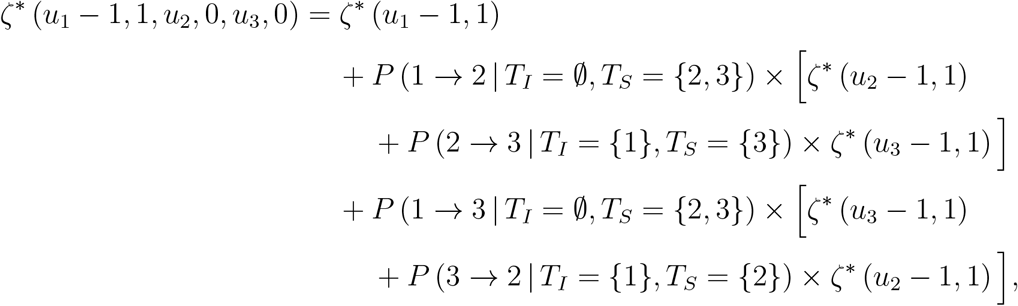

where the probabilities are defined as above for *P*(*k* → *j* | *T_I_, T_S_*).

### A.4 Modification to Keeling and Shattock’s Method

Keeling and Shattock [3] employed a deterministic model with a finite number of patches to determine the optimal allocation of vaccines to a metapopulation with the aim to minimise the final epidemic size. To calculate the final size for patch *ℓ*, for *ℓ* = 1,…, *m*, they used the following final size equation,

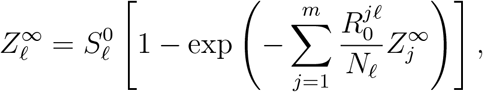

where 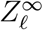 is the final size for patch *ℓ*, 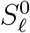 is the initial number of susceptible individuals in patch *ℓ*, *N_ℓ_* is the size of patch *ℓ* and 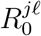 is the expected number of secondary cases produced in patch *ℓ* by a single infected individual in patch *j*, assuming all individuals in patch *ℓ* are initially susceptible (the notation is slightly modified from that used in Keeling and Shattock [3]). Then, the authors used simple recursion to attempt to solve the final size equation, given above, in the following way,

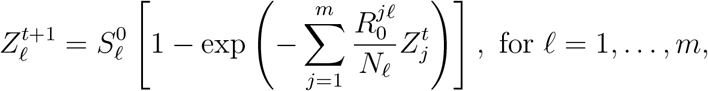

with the initial condition, 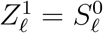, for *ℓ* = 1,…, *m*.

As we consider the seeding of infection into the population in our model, we adapt their method of calculation of the final epidemic size to suit our problem. To account for the patch in which infection is seeded (patch *k*), we modify the initial condition for the recursive method and also expand the final size equation to replace the first 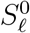 term with (*N_ℓ_* – *V_ℓ_*). The initial condition is 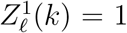, if infection is seeded in patch *k*, otherwise 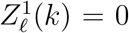. In a similar way, we set the second 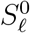 term in the expanded final size equation to be 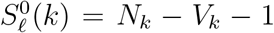, if infection is seeded in patch *k*, otherwise 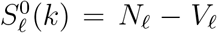. Although there is only a difference of one individual in 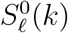 for the infected patch compared with the other patches, for small population sizes, this change can make a substantial difference. Another point of note is that we observed through numerical studies that there was little difference between Keeling and Shattock’s method and our revised method for reasonably large values of *R*_0_; however, as the value of *R*_0_ decreased, differences arose.

The revised final size equation used to determine the final size for patch *ℓ*, for *ℓ* = 1,…, *m*, where infection is seeded in patch *k*, is,

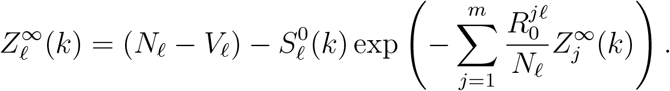

Using simple recursion, we can solve the revised final size equation by,

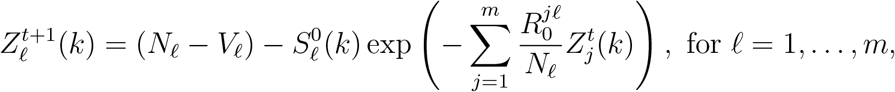

with

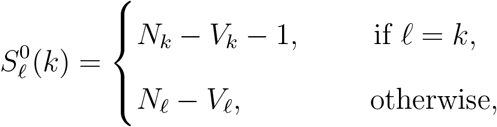

and initial condition,

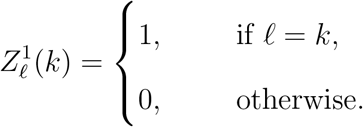

Another important point to note is the parameterisation used by Keeling and Shattock in their final size equation. For their model, they chose to use 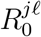 whilst our model uses *α_jℓ_β*, *β* and *γ*. Hence, we need a mapping between the two parameterisations:

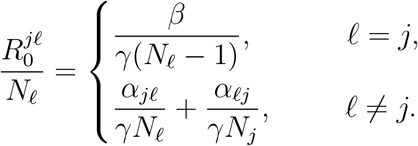

We have chosen this mapping as it corresponds to the transition rates of our model. Note that *ℓ* = *j* represents within-patch infection and we map this case to the within-patch component of the infection rate 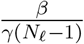. Similarly for *ℓ* ≠ *j*, we have cross-patch infection and so the cross-patch component of the infection rate 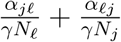.

As the recursive calculation of the final size for each patch, 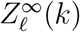, depends on the knowledge of which patch the infection has been seeded in, this tells us that the final epidemic size is dependent on the initial state. Hence, for a given initial state **u**(*k*) = (*u*_1_, 0,…, *u_k_* – 1,1,…, *u_m_*, 0), where infection has been seeded in patch *k*, the final epidemic size for **u**(*k*) is,

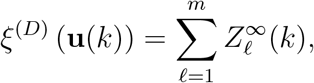

where 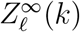 is the final size for patch *ℓ* obtained through recursion and is dependent on **u**(*k*) and hence the patch *k* where infection is seeded.

### A.5 Statistical testing procedure

As described in Section 4.4.2, when the optimal strategy and the mean final epidemic size cannot be obtained exactly due to the size of the problem, we are unable to use the relative difference in mean final epidemic size between the optimal strategy and the proposed strategy to determine how well the proposed strategy performs. Instead, to compare between the various proposed strategies (approximate, deterministic, equalising and fair), we simulate the mean final epidemic sizes for each proposed strategy and perform statistical tests to determine if the mean final epidemic size between the strategies are significantly different.

To simulate the mean final epidemic size, we perform many simulations using Sel-lke’s method [19] to obtain a set of final epidemic sizes for a given (*α, β*) pair and given strategy. Then, the statistical testing procedure used to compare the performance of our approximate strategy with the other strategies, deterministic, equalising and fair, consists of two statistical tests. The first is the one-way ANOVA test, which tests if the mean final epidemic size for all strategies are the same. The null and alternative hypotheses for this test are,

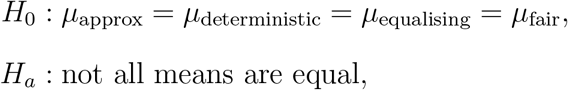

respectively, where *μ*_strategy_ is the mean final epidemic size for a given strategy. Using the `aov` function in R [20], we perform the one-way ANOVA test at a 5% significance level. This means that if the p-value of the test is less than 0.05, then we reject the null hypothesis, *H*_0_, and conclude that the mean final epidemic size for the strategies are not all equal.

As we are interested in determining if there is a significant difference in the mean final epidemic size between the strategies, if the null hypothesis for the one-way ANOVA test is not rejected, then there is insufficient evidence to conclude that the mean final epidemic sizes between the strategies are significantly different. However, if the null hypothesis is rejected, then not all the mean final epidemic sizes are the same and we are interested in which strategy or strategies are different from the approximate strategy. Hence, we make pairwise comparisons between the approximate strategy and the deterministic, equalising and fair strategies by performing Dunnett’s test [21, 22].

Dunnett’s test is a multiple comparisons test that tests various treatments with a control. This fits with our problem as the control is the approximate strategy and the other treatments are the deterministic, equalising and fair strategies. Hence, the null and alternative hypotheses are,

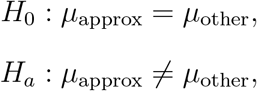

where *μ*_other_ is the mean final epidemic size for the other strategy (deterministic, equalising or fair). We use the `glht` function in R [23] to perform Dunnett’s test at a 5% significance level. Hence, if the p-value returned by the `glht` function is less than 0.05, then we reject the null hypothesis and conclude that there is a difference in the mean final epidemic size of the approximate strategy and the other strategy. Further, we can determine which strategy is better if we consider the difference in the mean final epidemic sizes. If the difference, *μ*_other_ – *μ*_approx_, is positive, then the approximate strategy is better than the other strategy. On the other hand, if the difference is negative, then the other strategy is better.

The other important aspect to consider when performing statistical tests is the underlying assumptions made. The assumptions of the one-way ANOVA test are:

- normality of observations in each group,
- equal variance of observations between groups,
- independence of observations within each group, and
- independence of observations between each group.

Dunnett’s test has the same assumptions of normality, constant variance and independence.

As each final epidemic size obtained from a single simulation is independent of final epidemic sizes obtained from other simulations, the independence assumption is satisfied. Next, we consider the assumption of constant variance. For this assumption to be reasonable, we require the following condition to be satisfied,

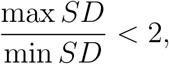

where min *SD* and max *SD* are the minimum and maximum sample standard deviations of the groups, respectively. For all problems considered, this assumption is satisfied except when the within-patch infection rate, *β*, is 0.5. Further, we also note from [24] and [25] that when the size of the groups are equal, moderate differences in the variance of each group does not have a serious impact. Hence, the assumption of constant variance is satisfied for our problem.

Lastly, we consider the normality assumption. For this assumption, we require the set of final epidemic sizes for each strategy of interest to be normally distributed. We know that this is not the case for the final epidemic size as it is either skewed or bi-modal. However, the central limit theorem tells us that for large enough samples, the sample mean is approximately normally distributed. As we perform 10^6^ simulations to obtain the set of final epidemic sizes, we can apply the central limit theorem to ensure that the sample means are approximately normally distributed [25]. Hence, for our problem, all assumptions are satisfied and it is appropriate for us to use the one-way ANOVA and Dunnett’s tests.

